# Robust and resource-optimal dynamic pattern formation of Min proteins *in vivo*

**DOI:** 10.1101/2023.08.15.553443

**Authors:** Ziyuan Ren, Henrik Weyer, Laeschkir Würthner, Dongyang Li, Cindy Sou, Daniel Villarreal, Erwin Frey, Suckjoon Jun

## Abstract

The Min system in *Escherichia coli* plays a crucial role in cellular reproduction by preventing minicell formation through pole-to-pole oscillations. Despite extensive research, predicting the onset of Min protein concentrations for oscillation and understanding the system’s robustness under physiological perturbations remains challenging. Our study aims to address these gaps. We show that the Min system’s dynamic pattern formation is robust across a wide range of Min protein levels and varying growth physiology. Using genetically engineered *E. coli* strains, we independently modulated the expression of *minCD* and *minE* in *E. coli* under both fast and slow growth conditions. This led to the construction of a MinD-MinE phase diagram, which revealed not just a large oscillation regime but also complex dynamic patterns such as traveling and standing waves. Interestingly, we found that the natural expression level of Min proteins is nearly optimal. Our work combines experimental findings with biophysical theory based on reaction-diffusion models, reproducing the experimental phase diagram and other key properties quantitatively. This includes the observation of an invariant wavelength of dynamic Min patterns across our phase diagram. Crucially, the success of our model depends on the switching of MinE between its latent and active states, indicating its essential role as a robustness module for Min oscillation *in vivo*. Our results underline the potential of integrating quantitative cell physiology and biophysical modeling in understanding the fundamental mechanisms controlling cell division machinery, offering insights applicable to other biological processes.

---

Pattern formation, a fundamental process in biological systems, enables organized development and functional differentiation. *Escherichia coli* offers an illustrative instance of this process, where the Min protein system displays pole-to-pole oscillations to aid symmetrical cell division (Fig. 1a). Despite the wealth of studies encompassing genetics^4–8^, structural biology^9–14^, biochemistry^15–22^, and biophysics^2,3,22–25^ dedicated to the Min system, several key questions remain unresolved.

**Fig. 1.**
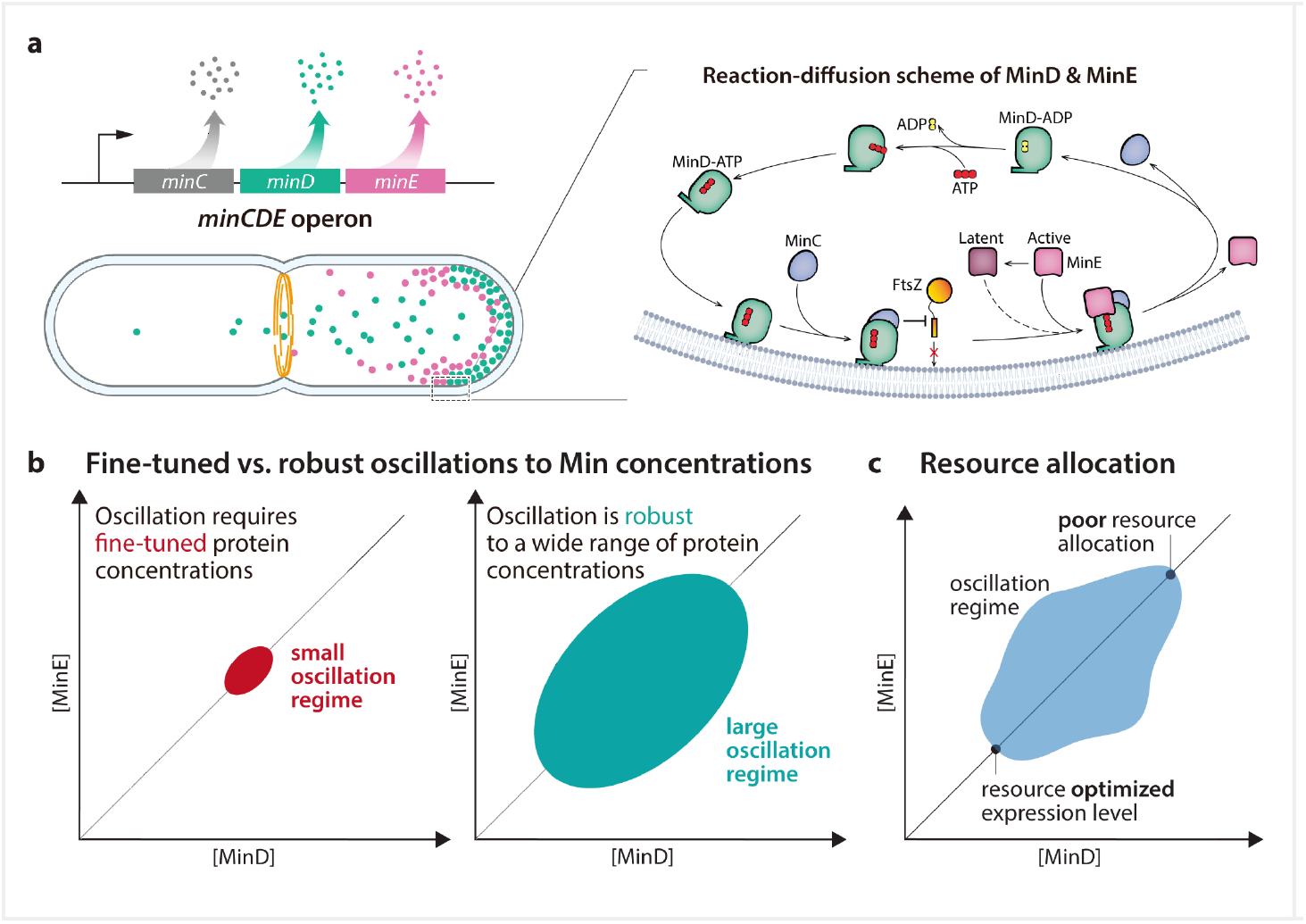
The Min system in the context of quantitative cell physiology. **a**, The *minCDE* genes are organized as an operon. The active form MinD-ATP binds to the membrane and recruits MinC, inhibiting the polymerization of the division protein FtsZ. MinE can switch from an active to a latent form^1^, and the membrane-bound MinD-ATP mainly recruits the active MinE. MinE stimulates ATP hydrolysis by MinD (i.e., MinD-ATP → MinD-ADP), and MinD-ADP is released to the cytoplasm. The pole-to-pole oscillations resulting from this biochemical reaction–diffusion system ensure that the concentration of MinD-ATP is highest at the cell poles, thus inhibiting cell division therein. **b**, So far, it is unknown at what Min protein concentrations pole-to-pole oscillations start, and whether the oscillation is sensitive (left) or robust (right) to physiological changes or perturbations to the protein concentrations. **c**, A related question is whether the production of Min proteins is resource optimal in wild-type.

Currently, our understanding is limited regarding how physiological perturbations affect Min oscillations. We broach the topic of the minimum Min protein concentrations for symmetric cell division (Fig. 1b), the expression levels in wild-type bacteria, and the possibility that these concentrations exemplify resource optimization (Fig. 1c). Additionally, *in vivo*, the genes encoding MinC, MinD, and MinE are expressed under a single operon (Fig. 1a). Yet, *in vitro*, pattern formation appears to withstand changes in the [MinE]:[MinD] ratio^1^. Does this presumed co-regulation impact successful cell division?

In this research, we scrutinize the robustness of the Min system’s dynamic pattern formation *in vivo*, considering variations in MinD and MinE concentrations, given that pole-to-pole oscillations hinge solely on the interactions between these proteins^26,27^. We constructed the first *in vivo* MinD-MinE phase diagram by creating unique *E. coli* strains that express *minCD* and min*E* in a gradient fashion. We conduct our experiments under both slow and rapid growth conditions, broadening the scope compared to previous works primarily centered on fast-growing *E. coli* cells^26,28–31^.

We augment our experimental study with biophysical theory, applying reaction-diffusion models that are grounded in the known biochemical reactions of the Min system^1–3^. Significantly extending the *in vitro* approaches so far, our *in vivo* and theory integration furnishes a holistic view of Min oscillations. We subsequently offer a minimal model that accurately interprets the experimental phase diagram, observed pattern types, oscillation periods, and wavelengths within a living cell.

Our integrative study sheds light on the role of the rapid conformational switch of MinE^1^, a central component in the Min system’s robustness. This ensures the system’s resilience against fluctuations in protein expression levels and growth conditions, as encountered by real cells in their natural environment. This results in a remarkably constant pattern wavelength across the entire phase diagram. Our phase diagram also indicates that, while co-regulation is not mandatory for successful proliferation, the wild-type protein expression closely approximates resource optimization.

## Construction and characterization of the *E. coli* strains for probing the MinD-MinE phase space

To explore the phase diagram of MinD-MinE interactions, we developed a variety of strains (Fig. 2; Materials and Methods). Initially, we created two tunable CRISPRi^32^ (tCRISPRi) strains capable of gradually repressing either the entire operon *minCDE* (SJ1696) or *minE* (SJ1697) (Fig. 2a). These repression strains allowed us to probe the MinD-MinE phase space in a diagonal manner {SJ1696 for [MinE] = [MinD]} or vertically {SJ1697 for [MinD] = [MinD]_wt_}, starting from the wild-type expression level (Fig. 2a).

**Fig. 2.**
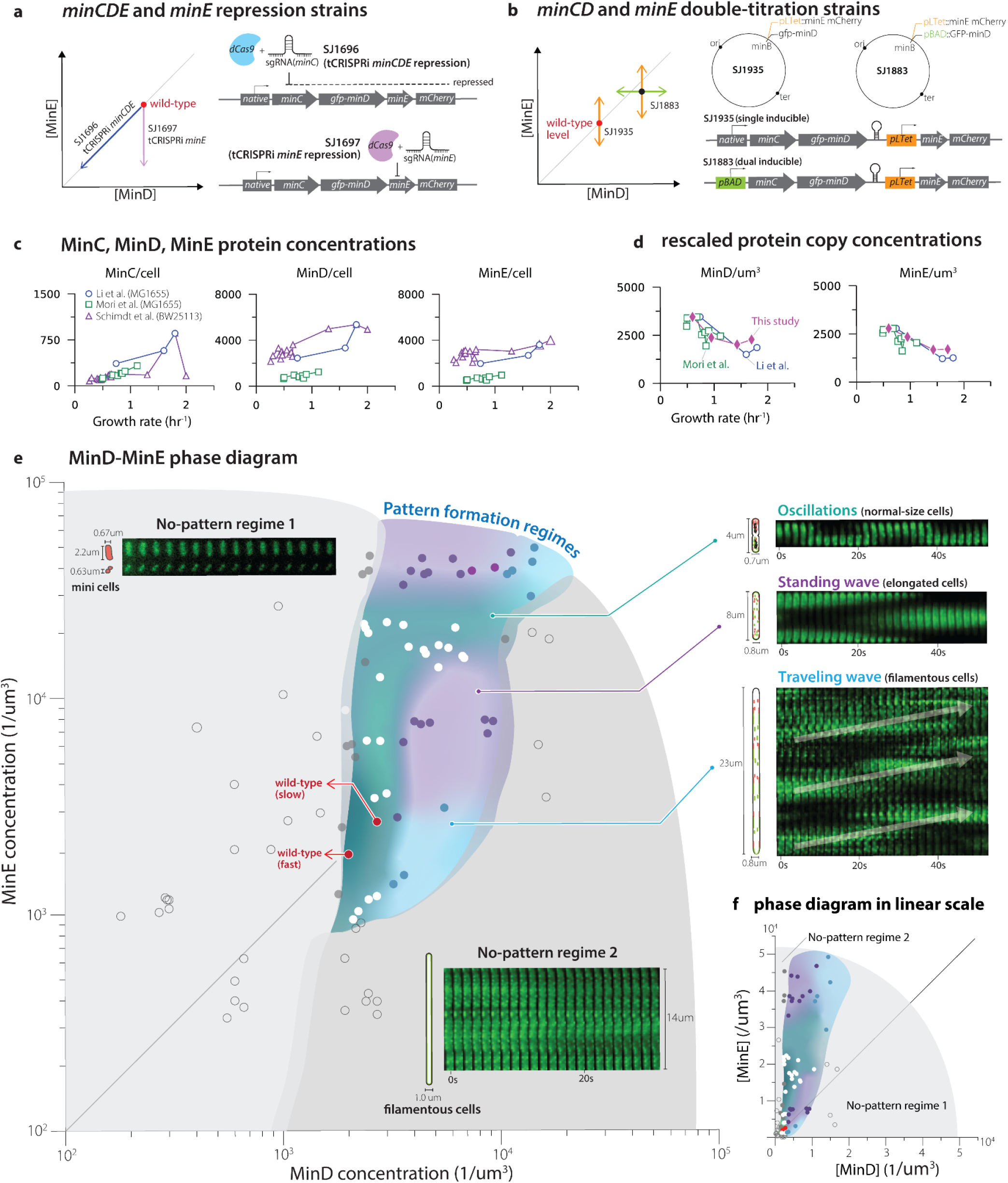
Exploring the [MinD]-[MinE] phase space using titratable strains, uncovering rich dynamic pattern formation *in vivo*. **a**, tCRISPRi strains for gradient repression of the whole *minCDE* (SJ1696) or *minE* (SJ1697). **b**, SJ1935: single inducible strain with pTet inducible promoter for gradient expression of *minE*. SJ1883: Dual inducible strain with arabinose-inducible promoter pBAD controlling the expression of *minCD* and tetracycline-inducible promoter pTet controlling the expression of *minE*, independently. **c**, MinC/D/E protein copy numbers per cell vs. growth rate measured in *E*.*coli* strain BW25113 (Schimdt et al.) and MG1655 (Li et al., Mori et al.). **d**, MinD/E protein concentration comparison between the published data and measurement in this study. The protein copy number per cell data (Fig. 2c) shows the opposite trend compared to the concentration data (**e**, experimentally-determined pattern formation phase diagram in the [MinD]-[MinE] space. Pattern formation regime contains wild-type oscillations (dark bluegreen) (SI Movie 1), traveling waves (blue) (SI Movie 2), and standing waves purple (SI Movie 3). For the two light-purple conditions, we observed transition patterns from standing to traveling waves. No patterns form if either the MinD and MinE concentrations are low or their ratio is severely disproportionate. In the regime without patterns, mini-cells form when [MinE]>>[MinD] (SI Movie 4), whereas filamentous cell forms when [MinE]<<[MinD] (SI Movie 5). Conditions that transition between pattern and no-pattern regimes are shown as gray data points, and these conditions show very weak non-static pattern. Simplified *in vivo* dynamics of MinD and MinE proteins are shown in the cartoon cells (red: MinE; green: MinD). (right) kymographs showing the representative dynamics of GFP-MinD *in vivo* in different regimes. Representative kymographs shown for each regime are obtained from the following conditions: Oscillation (20000/um^3 [MinE], 4000/um^3 [MinD]); standing wave (8000/um^3 [MinE], 8000/um^3 [MinD]); traveling wave (3000/um^3 [MinE], 6000/um^3[MinD]); no-pattern regime1 (10000/um^3 [MinE], 1000/um^3 [MinD]); no-pattern regime2 (300/um^3 [MinE], 2000/um^3 [MinD]) **f**, The same phase diagram is shown on a linear scale.

Furthermore, we constructed a dual-inducible strain (SJ1883) on the *E. coli* chromosome for independent, gradient expression of *minCD* and *minE* (Fig. 2b). Specifically, we replaced the native promoter with our previously developed pBAD* system ^32^ and inserted the pTet promoter between native *minD* and *minE*. For calibrating the pTet promoter under different growth conditions, we created an intermediate strain, SJ1935, which allowed for inducible expression of *minE* (Fig. 2b). Detailed dose-induction curves of the expression system can be found in the Supplementary Information Section 1. These *min*-inducible strains enabled us to explore the MinD-MinE phase space for more than a 10-fold expression range around the wild-type level.

To determine the Min protein concentrations, we employed fluorescence imaging and calculated the integrated fluorescence intensity normalized by the cell volume (Materials and Methods). To calibrate the fluorescence intensity to protein copy numbers, we extracted MinC, MinD, and MinE copy numbers from three published datasets based on mass-spectroscopy^33^, ribosome profiling, or Ribo-Seq, which quantifies protein levels by counting actively translating ribosomes^34^, as well as a hybrid method that combines both approaches^35^ (Fig. 2c). These published datasets reported a wide range of protein copy numbers per cell, with MinC ranging between 100-600, MinD between 1,000-5,000, and MinE between 500-4,000 (Fig. 2c). The observed variations can be attributed to differences in growth conditions, strains, and experimental methods. Among the three datasets, we compared our measurements with the Ribo-Seq and hybrid datasets. This choice was to ensure we could utilize our cell-size vs. growth rate data ^36^ to calculate the protein concentrations, because these two studies employed the same *E. coli* strain (MG1655) as ours (Fig. 2d). The trends of MinD and MinE concentration with respect to the growth rate were consistent among all three datasets and also aligned with our imaging-based measurements (Fig. 2d, Fig. S2b, and SI Section 2). Finally, to construct our phase diagram, we further calibrated our fluorescence intensity measurements using the Ribo-Seq data^34^, as their per-cell protein copy numbers agreed more closely with the mass spectrometry data (Fig. 2d & 2e, SI Sec. 2.2 Fig. S3).

We describe the strain construction and calibration in more detail in Materials and Methods and Supplementary Information section 2&3.

## The MinD-MinE phase diagram: oscillation is robust to physiological perturbations

With the four titratable strains we developed, we were able to explore the phase space of [MinD]-[MinE] by precisely controlling the expression levels of *minCDE* (SJ1696, SJ1883), *minCD* (SJ1883), and *minE* (SJ1697, SJ1883, SJ1935). This allowed us to probe a range of Min protein levels spanning about two orders of magnitude. To investigate the impact of growth physiology on Min pattern formation, we selected two growth conditions with average doubling times of 55 minutes (“slow”) and 25 minutes (“fast”) at 37°C. To construct the phase diagram and identify different regimes, we tracked the dynamics of GFP-MinD at various MinCDE levels using time-lapse microscopy and classified the observed dynamic patterns (Fig. 2e, right).

The phase diagram can be clearly divided into two regions based on whether cells under specific physiological conditions exhibit dynamic Min patterns. Strikingly, the pattern formation regime shows an asymmetric elongation in the [MinD]-[MinE] space (Fig. 2e and 2f), which was surprising as we initially expected oscillations to require a 1:1 MinD:MinE ratio based on the presumed co-regulation by the Min operon.

Within the pattern formation region, we identified multiple distinct regimes, including oscillations, standing waves, and traveling waves. Remarkably, we found that wild-type pole-to-pole oscillations remained robust across a wide range of Min concentrations, with approximately 5-6 fold changes in MinD concentrations, and 25-fold changes in MinE concentrations.

## The MinD-MinE phase diagram: the wildtype *minCDE* expression levels are near resource-optimal

Our phase diagram reveals that the wild-type expression levels of *minD* and *minE* are generally within a 2x range of the minimal protein concentrations required for oscillation, regardless of whether the cells were in slow- or fast-growth conditions. Given that the magnitude of gene expression noise typically falls within the range of 10% to 20% coefficient of variation (CV) ^38–40^, this shows that the Min protein levels at the wild-type expression levels are nearly resource-optimal for sustaining oscillations.

Even more so, constitutively expressed genes exhibit decreasing protein concentrations as the nutrient-imposed growth rate increases (Fig. 2d) ^37^. Consequently, the maximal achievable growth rate of *E. coli* sets the minimal protein concentrations occurring in wild-type cells growing under distinct conditions. Given that the concentration of MinD under fast-growth conditions is positioned at the onset of pattern formation, the expression of the Min operon appears to be strategically adjusted to be resource optimal but to allow Min oscillations under all growth conditions while also buffering fluctuations in protein concentrations. This finely tuned balance is in contrast to the large oscillation regime.

## The MinD-MinE phase diagram: oscillation is a subset of richer *in vivo* dynamic pattern formation

As we increased the expression levels of *minCD* or *minE*, we observed the emergence of dynamic patterns that go beyond what is seen in wild-type cells. Notably, we discovered two additional types of patterns: traveling waves and standing waves, both accompanied by the inhibition of cell division, resulting in filamentation.

Traveling waves exhibit a wavefront that moves in a unidirectional manner (Fig. 2d). In these waves, Min proteins continuously sweep through the entire cell, inhibiting septum formation on the cell membrane during cell growth and leading to the formation of filament-shaped cells. The specific locations of the traveling wave regimes on the phase plane depend on the MinE concentration, but both regimes appear at high MinD concentrations (Fig. 2c).

On the other hand, in standing waves, the GFP-MinD signal localizes periodically at fixed points within the cell (Fig. 2d). These patterns can be viewed as multiple neighboring pole-to-pole oscillations: Due to the defined wavelength of the Min oscillations (see below), the proteins oscillate between several “stripes” rather than between the poles. Although the Min proteins still exhibit the correct oscillation mode, cell division is suppressed, resulting in elongated cells. Similar to the traveling wave regimes, there are two distinct standing wave regimes separated by the oscillation regime: the first occurs along the diagonal line [MinD] = [MinE] but at 3x-10x the wild-type level, while the second occurs when [MinD] << [MinE].

## The MinD-MinE phase diagram: loss of pattern formation at high expression of *minD* or *minE*

When the expression levels of *minCD* and *minE* deviate significantly from the wild-type level, the dynamic patterns observed in the system eventually vanish. We identify two distinct “no pattern” regimes depending on the relative ratio of [MinE]/[MinD] (Fig. 2e). The origins of these regimes can be explained as follows:

In the regime where [MinE]/[MinD] >> 1, the presence of an excessive amount of MinE proteins strongly inhibits the binding of MinD to the cell membrane throughout the cell. Consequently, most of the MinD proteins remain in the cytoplasm and are unable to inhibit the formation of Z-rings at the cell poles, which are crucial for the division at mid-cell. As a result, mini cells are formed within this regime.

Conversely, in the regime where [MinE]/[MinD] << 1, the insufficient amount of MinE protein leads to the binding of MinD proteins all over the inner membrane, recruiting MinC from the cytoplasm. The excessive accumulation of MinD-MinC complexes on the inner membrane prevents the formation of the Z ring, resulting in filamentation of the cells.

In both cases, the disruption of the delicate balance between MinD and MinE proteins at extreme expression levels hinders the formation of the characteristic dynamic patterns.

## *In vivo* time-averaged GFP-MinD distribution shows narrow minimal concentration at mid-cell

In previous modeling studies, it was suggested that the distribution of Min proteins over time exhibits peaks at the poles of the cell, which corresponds to the role of the Min system in preventing the formation of minicells ^2,41^.However, until now, no systematic experimental study has compared the time-averaged concentration profile for different cell length during the cell cycle as well as under different growth conditions. Therefore, we conducted measurements to determine the *in vivo* time-averaged distribution of GFP-MinD under different growth conditions and at various cell lengths (Fig. 3).

**Fig. 3.**
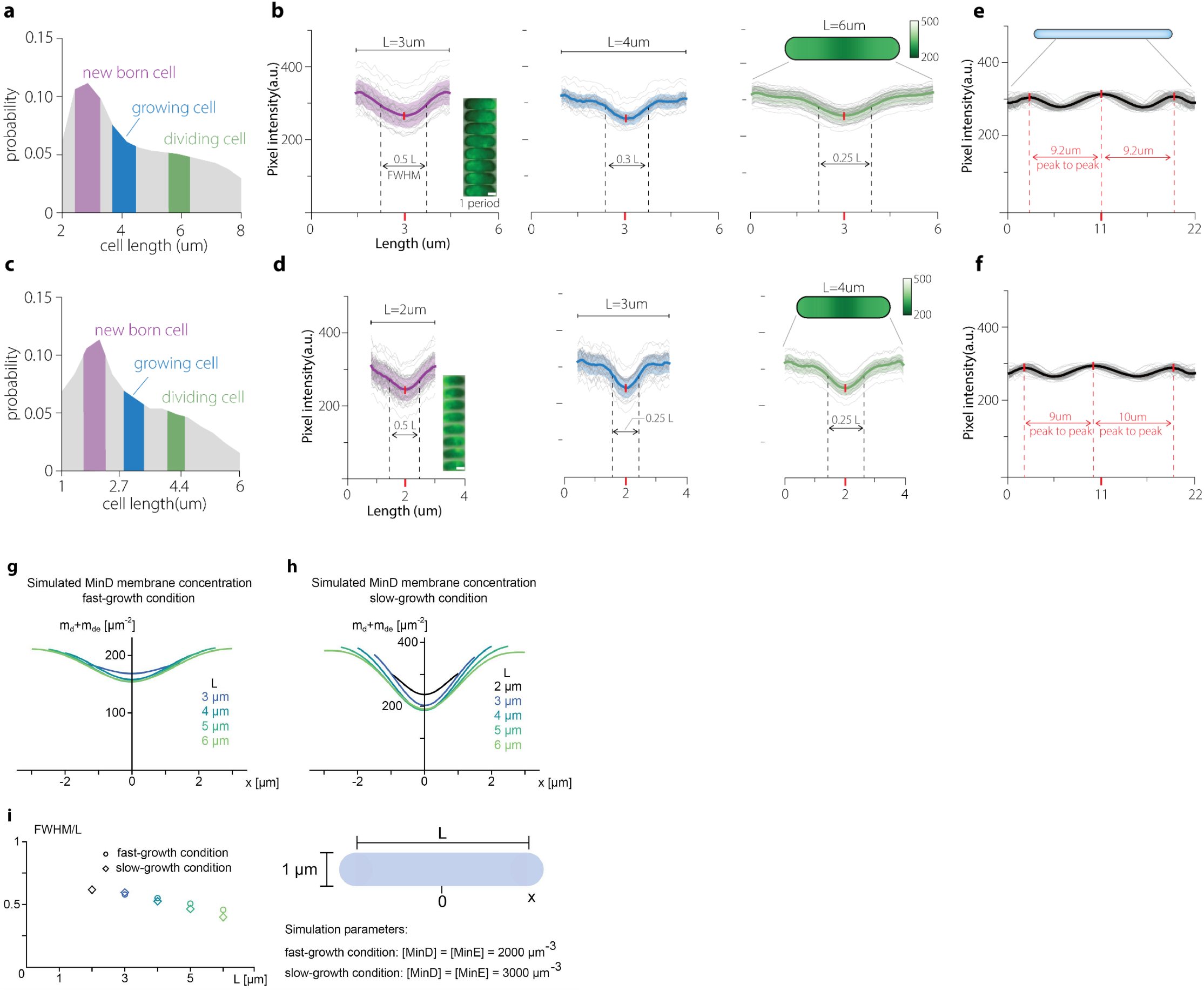
Time-averaged GFP-MinD distribution in wild-type cells and filamentous cells in fast and slow growth conditions. **a**, Length distribution of fast-growing wild-type cells. Sub-populations used for time-averaged GFP-MinD distribution analysis in panel **b** are labeled in purple, blue, and green. **b**, Time-averaged GFP-MinD intensity profile in fast growth conditions in short, intermediate, and long cells. We used the full-width-half-max (FWHM) as a measure of the intensity profile width. One full oscillation period was used for time-averaging (scale bar=1um in the kymograph). **c, d** same as **a, b** for slow growth conditions. The average cell size is smaller. **e, f** Time-averaged GFP-MinD distribution in filamentous (caused by over expression of minD) cells in fast (**e**) and slow (**f**) growth conditions. Only the cell center region was used for analysis, excluding the cell pole regions. **g, h** Simulated MinD membrane concentration at different cell lengths in fast (g) and slow (h) growth conditions. **i**, FWHM measured from plot (g) and (h). The Min protein concentrations are the only difference between fast- and slow-gorwth simulation, and the concentration values are chosen according to Fig. 2d.

Remarkably, the time-averaged GFP-MinD exhibited similar intensity profiles regardless of the cell’s growth physiology, prominently marking the mid-cell region (Fig. 3b,d). Intriguingly, the width of the minimum of the intensity profile at mid-cell also remains nearly constant during cell elongation, a feature that is also captured by the simulation of MinE-switch model using wild-type cell lengths (Fig.3g,h) Consequently, the relative width of the intensity distribution steadily decreases from 50% of the cell length to 25% from the experiment (Fig. 3b,d), but the decrease is less significant in the simulation (Fig. 3i). Although this is still significantly larger than the CV (approximately 5%) of the distribution of the septum position, it is comparable to the CV of other physiological parameters that we previously measured^39,42,43^. In other words, we expect the Min oscillation alone to be sufficient for cell viability, consistent with the mild phenotype observed in knockdown mutants of nucleoid occlusion genes ^44,45^.

Additionally, we induced filamentation by treating bacteria with cephalexin, a drug that inhibits Z-ring formation. The resulting elongated filamentous cells exhibited distinct wave-like patterns with a consistent wavelength of 9 µm -10 µm under both growth conditions (Fig. 3e,f). We will delve into further details regarding this observation in the next section.

## The crucial role of the MinE switch in experimental robustness

We describe the Min protein patterns using a reaction-diffusion model (Fig. 4a) based on proposed bimolecular reactions in the biochemical literature^1–3,13,46^. In this model, MinD-ATP, the active form of MinD, binds to the cell membrane and recruits additional MinD-ATP molecules. Membrane-bound MinD also recruits MinE, forming a MinDE complex where MinE stimulates the ATPase activity of MinD. Upon ATP hydrolysis, both inactive MinD (MinD-ADP) and MinE detach from the membrane^47–49^. In the cytosol, MinD-ADP is reactivated through nucleotide exchange.

**Fig. 4.**
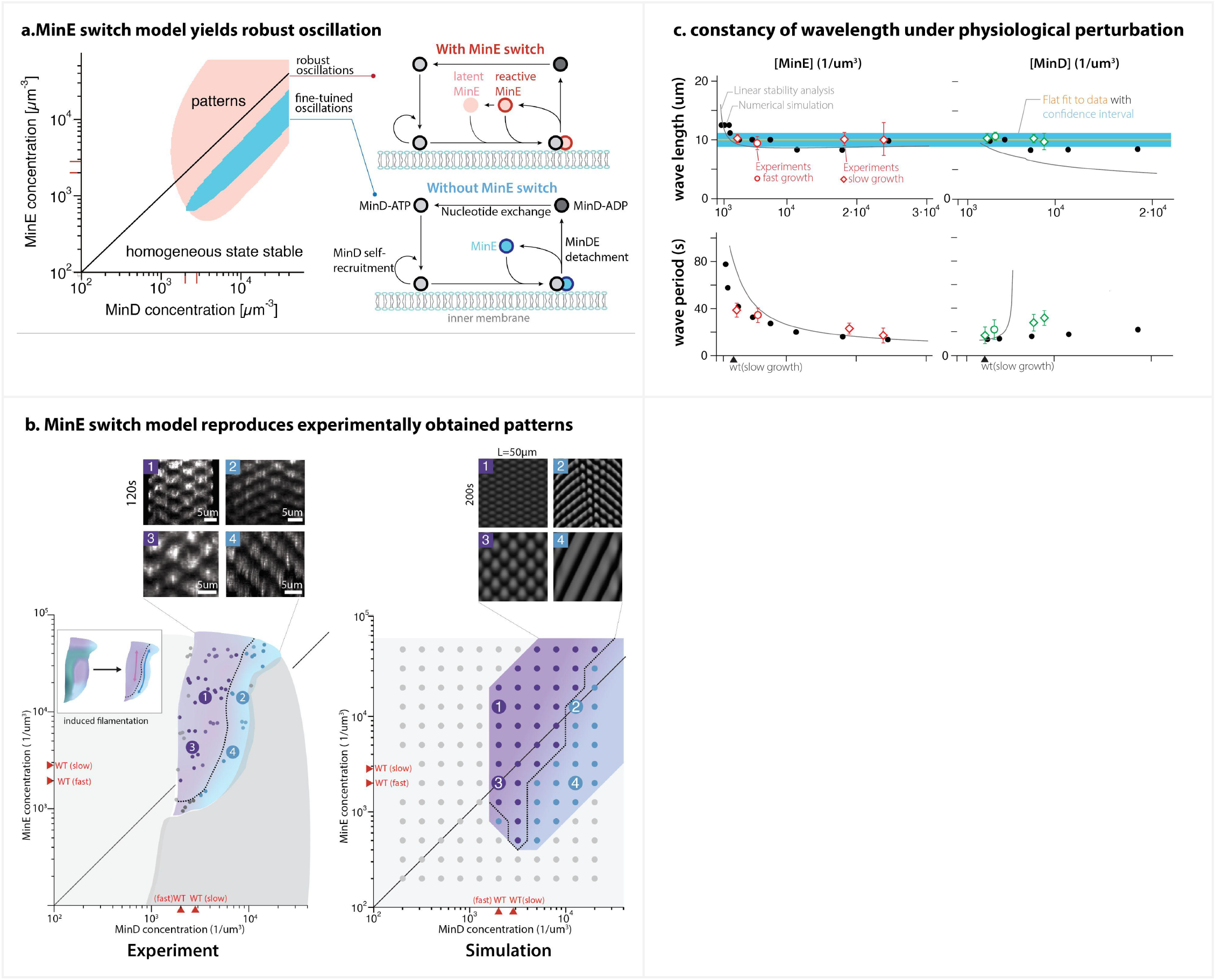
MinE-switch model and comparison between the experimentally obtained and the theoretically simulated phase diagrams. **a**, Comparison between the minimal and the MinE-switch model based on a linear stability analysis of the homogeneous steady state (see SI). Patterns grow out of small inhomogeneous perturbations of the uniform steady-state concentrations only within the shaded regions. The MinE-switch model features an additional latent MinE conformation that binds weakly to membrane-bound MinD-ATP. The resulting region of pattern formation covers a much larger area of the [MinD]-[MinE] phase space compared to the one predicted by the minimal model, which only permits pattern formation in a narrow diagonal stripe. **b**, After decoupling cell division from the Min protein expression levels, the experimentally-obtained patterns can be separated into either standing- or traveling-wave-type patterns (SI movies 6-9) with a gradual crossover between both. The experimental phase diagram compares well with the phase diagram obtained with the switch model (see Methods). Representative kymographs at similar [MinD] and [MinE] levels from both experiments and simulation are shown at the top. **c**, Wavelength and period of the patterns obtained from experiments and simulation. The Min protein pattern formation shows wavelength constancy (∼10um) over a large range of Min protein concentrations (top). The wave period decreases with increasing [MinE] and increases with increasing [MinD] (bottom). The simulations are performed at a fixed MinD concentration of 3000 µm^-3^ or MinE concentration of 27000 µm^-3^, respectively.

Additionally, we incorporate the conformational switch of MinE between its reactive state and latent state induced by MinD^1,13^. In the reactive state, MinE readily binds to the inner membrane (Fig. 4a). Conversely, in the latent state, MinE has a weak affinity for the membrane as its interaction sequence with MinD is conformationally buried. The minimal network without the MinE switch already qualitatively reproduces pole-to-pole oscillations observed in wild-type cells, as well as the dependence of dynamic patterns on cell geometry^3,50^. However, linear stability analysis of the homogeneous protein distribution reveals that patterns only form within a narrow range of the [MinE]/[MinD] ratio (Fig. 4a; Ref ^3^), in clear contrast to our experimental observations (Fig. 2e).

The model incorporating the MinE switch explains the robustness of pattern formation observed in *in vitro* experiments^1^. The underlying mechanism lies in the latent state acting as a reservoir that buffers high concentrations of MinE and enables pattern formation even at high [MinE]/[MinD] ratios. We simulated this “switch model” within the cellular geometry. As depicted in Fig. 4a, the switch model adequately accounts for the robustness of pattern formation in vivo, spanning over one order of magnitude in MinE concentration. The pattern-forming regime extends to higher MinD concentrations in the model compared to the experiments, which we attribute to the aggregation of MinD not considered in the model. (see SI section 6).

## The MinE-switch model captures the various types of patterns observed in filamentous cells

The success of the switch model in reproducing the robustness of pattern formation prompted us to investigate whether it also explains the pattern types observed in the experimental phase diagram (Fig. 2e). The phase diagram reveals that Min protein patterns are influenced by two interconnected processes: (1) the formation of different pattern types by the Min proteins and (2) the length at which cells divide, which in turn can impact the pattern types observed. To isolate pattern formation from cell size, we re-explored the phase diagram, treating cells with cephalexin to suppress cell division in those regions of the phase diagram where cells do not grow filamentous. As a result, the cells grow filamentous, and the Min patterns transition into either standing-wave-type patterns at a high [MinE]/[MinD] ratio or traveling-wave-type patterns at a low [MinE]/[MinD] ratio (Fig. 4b). Previous studies by Raskin and de Boer^26^ also observed standing-wave-like behavior of MinD in FtsZ-filamentous cells. The arrangement of the phase diagram in our study is similar to the behavior observed in recent *in vitro* experiments on pole-to-pole oscillations and circular traveling-wave patterns in spherical microdroplets^51^, but opposite to observations on supported lipid bilayers even at low bulk height^52^.

Remarkably, our simulations using the switch model closely reproduce the pattern formation regimes observed in filamentous cells. We obtain standing-wave patterns at high [MinE] and traveling-wave-type patterns at low [MinE] (see Methods for the model parameters). The transition between the two regimes is gradual, consistent with experimental observations (Fig. 4b). The close resemblance between the experimental and modeling phase diagrams (Fig. 4b) supports the notion that the MinE-switch model encompasses the primary interactions governing Min pattern formation *in vivo*.

While the MinE switch enables robust pattern formation across a wide range of model parameters, the arrangement of distinct pattern types in the phase diagram imposes constraints on parameter selection (see Methods/SI). Our combined analysis reveals that the observed pattern-forming behavior arises with a significantly reduced self-recruitment of MinD onto the membrane compared to the parameters inferred from the minimal model without the MinE switch.^1,3^ With the inclusion of the MinE switch, less strong positive feedback in the MinD attachment is sufficient to explain the formation of standing and traveling waves.

## Oscillation period of dynamic Min patterns

We also quantified the oscillation period and wavelength of the patterns. Measurement in the filamentous cells shows that the oscillation period decreases with increasing MinE concentration, and increases with increasing MinD concentration (Fig. 4c). The behavior is quantitatively reproduced in the simulations with the MinE-switch model (black circles in Fig. 4c). The oscillation period increases with decreasing MinE:MinD concentration ratio because the reduced (relative) amount of MinE decreases the rate with which sufficient MinE is recruited to a MinD domain to induce detachment. Similarly as for the reduced region of pattern formation (Fig. 4a, b), we attribute the increased experimental oscillation period at high MinD concentrations compared to the simulations to the aggregation of MinD and a thereby reduced detachment rate of MinD from the membrane.

## The wavelength of dynamic Min patterns exhibits remarkable invariance

Throughout the entire phase diagram, the pattern wavelength remains remarkably invariant, regardless of variations in MinD and MinE concentrations or growth conditions. The wavelength consistently maintains a value of approximately 10µm (as shown in Fig. 4c). This observation is supported by the numerical analysis of the switch model, which also demonstrates the invariance of the wavelength. Notably, when close to the onset of pattern formation at low MinD concentration, the wavelength and oscillation period can be determined through linear stability analysis due to the supercritical nature of the onset^53^ (indicated by the black lines in Fig. 4c). More detailed information can be found in the supplementary information provided.

## Conclusion and perspective

In this study, we have made substantial advancements in understanding a paradigmatic cell division control mechanism in bacteria. Using *E. coli* as our model organism, our study offers new insights into how the Min protein system’s pattern-forming behavior intertwines with cellular physiology. To study this link systematically, we construct the first *in vivo* MinD-MinE phase diagram, addressing long-standing questions on the efficiency of wild-type expression levels, the importance of *minCDE* co-regulation, and the robustness of Min oscillations. Our findings underscore the resiliency of pole-to-pole oscillations across varying MinD and MinE protein concentrations and, at the same time, indicate that the physiological levels of MinD and MinE align closely with resource optimization, highlighting the system’s efficiency. Moreover, the phase diagram unveiled the emergence of different growth phenotypes and other dynamic patterns, including standing and traveling waves, *in vivo*.

We show that a minimal reaction-diffusion model based on the MinD-dependent conformational switch of MinE not only captures the various regimes of the MinD-MinE phase diagram but also provides quantitative predictions for critical observables like the oscillation period and pattern wavelength. This congruence between model predictions and experimental data underlines the pivotal role of MinE switching between a latent and a reactive conformation. Without the latent state, the model predicts oscillations only within a finely-tuned range of *minCDE* expression. Our analysis affirms that the MinE switch is a crucial robustness mechanism for Min oscillation *in vivo*, providing mechanistic insight into the experimentally observed cell-division behavior.

Another intriguing finding from our combined study is the invariant wavelength of Min patterns around 10 µm, irrespective of *minCDE* expression levels and growth physiology. This invariance provides additional robustness to symmetric cell-division in *E. coli*, as it can determine the mid-cell. Indeed, studying the evolution of the time-averaged MinD concentration profile in wild-type cells during the cell cycle, we show that the fixed, intrinsic pattern wavelength allows for a sharp concentration minimum at midplane that does not widen as the bacteria grow. Therefore, as long as the average dividing cell is shorter than 10 µm, the mid-cell position is robust. Intringuingly, the average size of fastest-growing (thus largest) *E. coli* cells at division is approximately 8 µm long^42^. Furthermore, while a linear stability analysis well accounts for the wavelength of weakly nonlinear patterns, understanding the basis of wavelength selection in the genuinely nonlinear regime is a critical open question. Moreover, the invariance of the Min wavelength could have other biological implications, such as cell-size control^54,55^, which is an exciting avenue for further research.

In a broader context, bacteria utilize a plethora of mechanisms for spatial cell division control. We have identified the MinE switch as a robustness module in *E. coli*’s cell division control via Min oscillations. Since accurate cell division is pivotal for proliferation, it is essential to explore molecular mechanisms that ensure robust spatial control in bacteria employing different strategies. Moreover, there is a fundamental balance between resource efficiency and robustness against fluctuating gene expression levels. Comprehending the interplay between different spatial control strategies, their robustness, and protein-concentration homeostasis will be key to understanding the accurate selection of division sites across generations under varying conditions.

## Materials and Methods

### Strains used for the MinD-MinE phase diagram

To measure the dynamic Min patterns and quantify the Min protein concentrations using imaging, we N-terminally fused *sfGFP* to *minD* and inserted a transcriptional reporter *mCherry* downstream of *minE*. Thereby, fused *sfGFP* enables us to monitor the Min patterns while MinE remains unmodified. The latter is necessary since the fluorescent fusion of MinE proteins is not fully functional^56,57^. All experiments were performed under the same imaging condition.

All our strains used in this work are derived from MG1655^58^. SJ1695 has an N-terminus fusion of sfGFP^59^ to *minD* and mCherry^60^ as a transcriptional reporter encoded downstream of *minE* in our tunable CRISPR interference (tCRISPRi) strain without the sgRNA.^61^ Construction of the gradient repression strains SJ1696 and SJ1697 is based on SJ1695 with sgRNA sequences targeting *minC* (SJ1696) and *minE* (SJ1697).

We constructed the *minE* inducible strain SJ1935 from SJ1695 by inserting an inducible promoter pTet between *minD* and *minE*, the *rrnB* terminator between *minD* and *pTet*, and the repressor expression system pTet::tetR at the yfdG locus. We subsequently built the dual inducible strain SJ1883 by replacing the native promoter of the Min operon with the pBAD^39^ promoter.

| Strain | Parent strain | Genotype |
| --- | --- | --- |
| SJ1695 | SJ797 (MG1655 with pBAD<sup>32</sup> system) | pBAD_dCas9 ΔLacI ΔaraE araFGH<>spec lacYA177C; galM// tet-sacB//gmpA; minC-GFP-minD-minE-SD-mCherry pSIM18 |
| SJ1696 | SJ1695 (recombination to replace tet-sacB segment with minC sgRNA) | pBAD_dCas9 ΔLacI ΔaraE araFGH<>spec lacYA177C; galM// minC sgRNA//gmpA; minC-GFP-minD-minE-SD-mCherry pSIM18 |
| SJ1697 | SJ1695(recombination to replace tet-sacB segment with minE sgRNA) | pBAD_dCas9 ΔLacI ΔaraE araFGH<>spec lacYA177C; galM// minE sgRNA//gmpA; minC-GFP-minD-minE-SD-mCherry pSIM18 |
| SJ1883 | SJ1882(recombination to remove araBAD gene for pBAD system) | pBAD::minC GFP-minD KmR pTet::minE mCherry; araC cat ΔaraBAD; ΔsgRNA; yfdG pTet::tetR |
| SJ1935 | SJ1934 (recombination to insert tetR gene for pLTet system) | minC GFP-minD KmR; pTet::minE mCherry; Ptet::tetR<>yfdG ΔaraBAD; ΔsgRNA; |

### Growth conditions

Strains kept at -80C glycerol stocks were immediately streaked on an LB medium plate with proper antibiotics and inducer concentrations [SJ1935 with chlortetracycline (cTc) and SJ1883 with both cTc and arabinose] for recovery at the wild-type Min protein levels. Specifically, 50ug/ml spectinomycin for SJ1695, SJ1696, and SJ1697; 50ug/ml kanamycin+80ng/ml cTc for SJ1935; and 50ug/ml kanamycin+25ug/ml chloramphenicol+80ng/ml cTc+ 0.05%(w/v) arabinose for strain SJ1883.

Single colonies were picked and inoculated into 1ml LB pre-culture for overnight growth in the 37C water bath in the same conditions as described above for the LB nutrient plate. For the fast-growth condition experiments, we back-diluted the overnight culture 1:1000 into MOPS-rich 0.2% glycerol with the same amount of antibiotics added. Gradient expression of *minCDE* was achieved by adding differential amounts of inducers after 1:1000 back dilution in strains SJ1883 (both arabinose and cTc) or SJ1935 (cTc only). Gradient repression of *minCDE* or *minE* was achieved by adding differential amounts of arabinose after 1:1000 back dilution in strain SJ1696 or SJ1697. For the slow-growth experiments, there was a considerable lag period between the back dilution and the start of growth when using a 1:1000 dilution. Therefore, we first back-diluted at a lower 1:100 ratio and allowed the cells to grow before the second 1:10 back-dilution. For each back-dilution step, the procedure is the same as we did in the fast-growth experiments. All experiments were performed at 37C.

### Imaging sample preparation

We started preparing for imaging when the back-diluted cells reached OD600 = 0.3. We used a growth-medium-based 2% agarose (Sigma A9539-500G) pad on a Wilco dish (WillCo-dish® KIT-5040), which was pre-warmed in the 37C warm chamber before we put it on the agar pads. On the Willco dish, we prepared different agarose pads by varying the inducer concentrations to match the inducer concentrations in the liquid cultures (except no antibiotics were added for agar-pad imaging). The perimeter of the coverslip was sealed with dielectric grease to prevent the agar pad from drying during imaging at 37C.

### Imaging and Microscopy

Fluorescence image acquisition was conducted on a Nikon Ti-E inverted microscope using the Prime95B sCMOS camera. We used Coherent OBIS 488nm (100mW) and 561nm (50mW) lasers at 100% power with 200ms exposure time to excite sfGFP and mCherry, respectively. The sfGFP was imaged every 2 seconds to capture the dynamics of MinD proteins. For obtaining the statistics of fluorescence signals, we imaged both sfGFP and mCherry every 5 minutes with multiple fields of view. On average, we analyzed 500-2000 cells for each inducer condition for fluorescence intensity analysis.

### Image processing and data analysis

We used Delta2.0^62^ for cell segmentation and tracking during cell growth on the agar pads. After cell segmentation, we quantified the protein concentration using the integrated total fluorescence intensity normalized by the cell volume. We computed the volume by approximating the cell as a cylinder with two hemispherical caps.

### Reaction–diffusion model

We model the geometry of filamentous cells by a spherocylinder composed of a cylinder of length *L* = 50 µ*m* and radius *R* = 0. 5 µ*m* and two hemispherical caps at its ends with the same radius. The interior of the spherocylinder represents the cytosol. In the switch model, the cytosol contains the inactive MinD-ADP, active MinD-ATP, reactive MinE, and latent MinE states (Fig. 4a). The evolution of their concentration fields 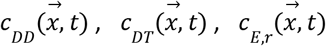, and 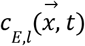, respectively, is modeled by the reaction–diffusion equations

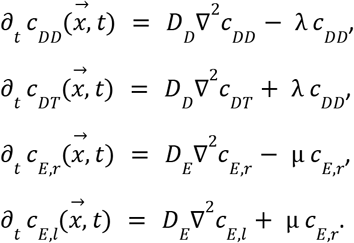

The diffusion coefficients and reaction rates are given and explained in Table 2. The surface of the spherocylinder represents the inner cell membrane. The surface concentration fields of membrane-bound MinD proteins 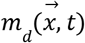 and MinDE complexes 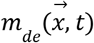 follow the equations

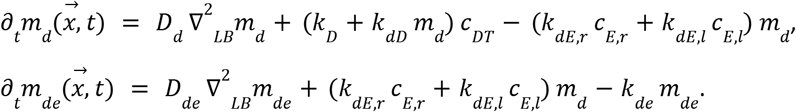

Here, the Laplace-Beltrami operator ∇^2^ _*LB*_ is used to describe diffusion along the (curved) membrane. For the rate constants, we refer again to Table 2. The attachment of proteins onto the membrane implies a depletion of the cytosolic protein concentrations. This interaction is described by the reactive boundary conditions

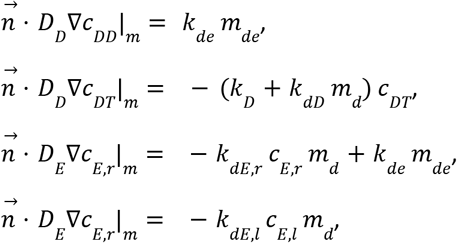

Where 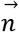 denotes the outward-pointing unit normal vector of the surface, and · |_m_ denotes the evaluation on the membrane, i.e., on the surface of the spherocylinder.

**Table 2:** Parameters of the switch model employed in the simulations in Fig. 4.

| Parameter | Value | Description |
| --- | --- | --- |
| $D_D$ | $16 \mu m^2 s^{-1}$ | MinD cytosolic diffusion coefficient |
| $D_E$ | $10 \mu m^2 s^{-1}$ | MinE cytosolic diffusion coefficient |
| $D_d$ | $0.05 \mu m^2 s^{-1}$ | MinD membrane diffusion coefficient |
| $D_{de}$ | $0.05 \mu m^2 s^{-1}$ | MinDE membrane diffusion coefficient |
| $k_D$ | $0.3 \mu m s^{-1}$ | MinD membrane attachment rate |
| $k_{dD}$ | $7.5 \cdot 10^{-4} \mu m^3 s^{-1}$ | MinD self-recruitment rate |
| $k_{dE,r}$ | $0.75 \mu m^3 s^{-1}$ | Recruitment rate of reactive MinE |
| $k_{dE,l}$ | $5 \cdot 10^{-6} \mu m^3 s^{-1}$ | Recruitment rate of latent MinE |
| $k_{de}$ | $1 s^{-1}$ | Hydrolysis / MinDE membrane detachment rate |
| $\lambda$ | $5 s^{-1}$ | Nucleotide exchange rate |
| $\mu$ | $20 s^{-1}$ | MinE switch rate |
| $\lambda$ | $5 s^{-1}$ | Nucleotide-exchange rate |
| $\bar{\rho}_D$ | Varied [$\mu m^{-3}$] | Total average MinD concentration |
| $\bar{\rho}_E$ | Varied [$\mu m^{-3}$] | Total average MinE concentration |
| $R$ | $0.5 \mu m$ | Radius of spherocylinder |
| $L$ | $50 \mu m$ | Length of the cylindrical part of the spherocylinder |

The minimal model without the MinE switch (Halatek, Frey, Cell Reports, 2012) is recovered from the above equations by summing up the reactive and latent MinE states *c*_*E*_ = *c*_*E,r*_ + *c*_*E,l*_and setting the recruitment rates for both states to the same value *k*_*dE,r*_ = *k*_*dE,l*_ = *k*_*dE*_.

Due to the small radius of the spherocylinder, the dynamics can be approximated by a reduced one-dimensional model that assumes the cytosolic concentration as constant within a crosssection of the spherocylinder (see SI). The dynamics of the concentration fields integrated over each crosssection evolve on the one-dimensional line, and we model the cell poles as no-flux boundary conditions constraining the evolution to an interval of length *L* = 50 µ*m*. This reduced dynamics is used to perform an extensive study of the rate constants in the switch model. Moreover, the reduced dynamics on the infinite line is used to perform the linear stability analysis (Fig. 4a, black lines in Fig. 4c, and SI section 8).

### Parameter choice

The reaction parameters employed in Ref.(Denk et al. 2018) for the switch model give rise to traveling waves in most of the phase diagram in contrast to the experimentally observed standing- and traveling-wave regions (see SI ref. To Fig. S11). The parameter set used to describe the experimentally determined phase diagram is given in Table 2. It results from a broad parameter study because the reaction rates are unknown experimentally (see SI section 10.2).

We use the cytosolic diffusion coefficients of MinD and MinE that have been determined experimentally *in vivo* using fluorescence correlation spectroscopy(Meacci et al. 2006). Estimates for their membrane diffusion constants have been obtained in *in vitro* measurements(Loose et al. 2011). Because the cell membrane of live bacteria is crowded by diverse molecules, we choose low membrane diffusion constants comparable to the values measured *in vitro* at high MinD densities on the membrane. We increased the value compared to the value used in the minimal model in Ref.(Halatek and Frey 2012) to reproduce the experimentally determined pattern wavelength better. However, the wavelength only changes weakly with the membrane diffusion constant (data not shown).

The nucleotide exchange rate λ = 5 *s*^−1^ is chosen to meet the lower bound determined in Ref.(Meacci et al. 2006). The other reaction rates are determined to reproduce the experimental phase diagram. To match the overall region of pattern formation, following Denk et al.(Denk et al. 2018), a small value is chosen for the recruitment rate *k*_dE,l_ of the latent MinE state compared to the rate *k*_dE,r_ for the reactive MinE state. This allows the formation of a MinE reservoir. The switching rate µ = 20 *s*^−1^ is chosen to describe a fast switch. It is reduced compared to the value µ = 100 *s*^−1^ employed in Ref.(Denk et al. 2018) to enlarge the phase-space region showing standing-wave patterns (see SI, Fig. S17). The hydrolysis rate *k*_de_ adjusts the threshold of pattern formation at low total MinE concentrations, and it is fixed to match the experimentally observed oscillation frequency of the Min patterns (see SI, Fig. S19). The linear MinD attachment rate *k*_D_ influences the onset of pattern formation at low total MinD and MinE concentrations. The MinD self-recruitment rate *k*_dD_ affects the pattern threshold at low total MinD concentrations and the pattern wavelength. Both *k*_D_ and *k*_dD_ are then fixed such that the pattern types and wavelengths are matched in the phase diagram (see SI, Figs. S16, S18).

For the minimal model without the MinE switch, we use the parameters determined in Ref. (Halatek and Frey 2012), evaluating the hydrolysis rate *k*_de_ at 37°C. We scale the nonlinear reaction rates *k*_dD_ and *k*_dE_ by a factor of 1/60 to rescale the protein concentrations by a factor of 60 and move the region of instability to concentration values comparable with the experimental phase diagram (cf. Fig. 4a).

### Numerical simulation

The spherocylindrical simulation domain is cylindrically symmetric. Moreover, the observed patterns show the same symmetry after an initial transient. Employing this symmetry, we simulate the reaction–diffusion dynamics in a two-dimensional radial slice of the spherocylinder for *T* = 3000 *s* using COMSOL Multiphysics 6.0 & 6.1 (see SI). The simulations employ a finite-element discretization on a triangular Delaunay mesh with linear Lagrange elements for the cytosol (bulk) and quadratic elements on the membrane (boundary). Initially, the total protein mass is homogeneously distributed in the cytosolic states *c*_DT_ and *c*_E_ or *c*_E,l_ with weak random perturbations (uniformly distributed within ± 1% around the homogeneous concentration). The reduced one-dimensional dynamics is solved using a uniform finite-differences discretization implemented in Mathematica 13.0 & 13.1 using second-order central differences. Again, the dynamics is simulated for *T* = 3000 *s*. In all simulations, we employ the freedom in the choice of units, and the fast-growth wild-type concentrations are normalized to average MinD and MinE concentrations 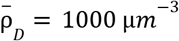 and 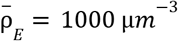 by scaling the nonlinear rates 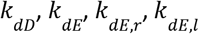 by a factor 2.

Mathematica 13.0 & 13.1 is also used to perform the linear stability analysis and analyze the numerical simulations. The classification into standing- and traveling-wave patterns is based on the kymographs of the total density of membrane-bound MinD. Traveling waves give rise to diagonal lines of high (low) density, while standing waves give rise to disconnected domains of high or low MinD membrane density (cf. Fig. 4b). To distinguish both pattern types, the number of high- and low-density domains per oscillation period and wavelength is counted (see SI). The threshold corresponds to a pattern showing a traveling wave in approximately half of the cell and a standing wave in the opposite half. The oscillation period and pattern wavelength are determined by Fourier analysis. The position of the maximum of the spatially averaged power spectrum and temporally averaged structure factor are used as measures for the period and wavelength, respectively (see SI).

Exemplary Comsol setup files and the Mathematica notebooks are available at https://github.com/henrikweyer/Min-in-vivo.

