## Supplementary information for "Robust and resource-optimal dynamic pattern formation of Min proteins *in vivo*"

**4 Cephalexin treatment: decouple cell length from the Min protein levels 6**

**5 The Min system shows similar oscillation period at similar Min protein expression levels in different growth conditions 11**

**6 Comparison between the experiment and theoretically predicted pattern formation regime 11**

**7 Models for the *E. coli* Min system in cellular geometry 13**

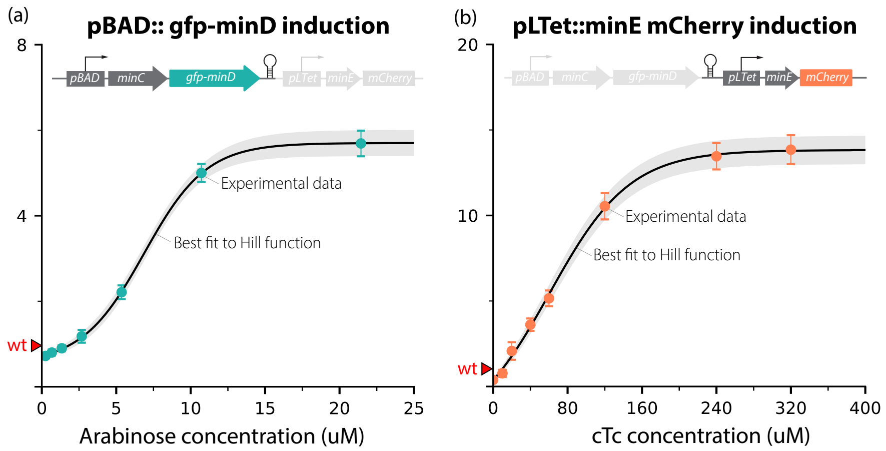

Figure S1. Dose-induction of pBAD::gfp-minD and pLTet::minE mCherry levels normalized by the wild-type level. (a) dose-induction curve of pBAD::gfp-minD:*in vivo* GFP-MinD expression level increases from below the wild-type level to roughly 6-fold of the wild-type level as the arabinose concentration increases. (b) dose-induction curve of pLTet::minE mCherry:*in vivo* MinE mCherry expression level increases from below the wild-type level to roughly 13-fold of the wild-type level as the cTc concentration increases.

### 1 Dose induction curve for pBAD::gfp-minD and pLTet::minE mCherry

Gradient control of the Min protein expression levels under inducible promoters requires the information between the amount of inducers added and the resulting concentrations. Here in Figure S1 we showed the dose induction curves of the two inducible systems, pBAD and pLTet, used for generating the experimental phase diagram. Normalized GFP and mCherry voxel intensities (wild-type levels are 1 on the plot) within the cell were used as proxies of the expression levels of minC/D and minE, respectively. Both inducible systems were able to achieve gradient expression of the Min proteins from below the wild-type levels to much higher than the wild-type levels [see Fig. S1].

### 2 Min protein concentration calibration

#### 2.1 Min protein copy number from published data

Currently, there are three sets of proteomic data that we find relatively reliable. The first is the ribosome profiling data (Ribo-Seq) Ref. [1]. The second is mass spectroscopy Ref. [2] . The third is a hybrid version of ribosome profiling and mass spectroscopy Ref. [3]. We also looked for the copy number and concentration information in Sunney Xie's classic paper Ref. [4], which used similar approaches. Unfortunately, they did

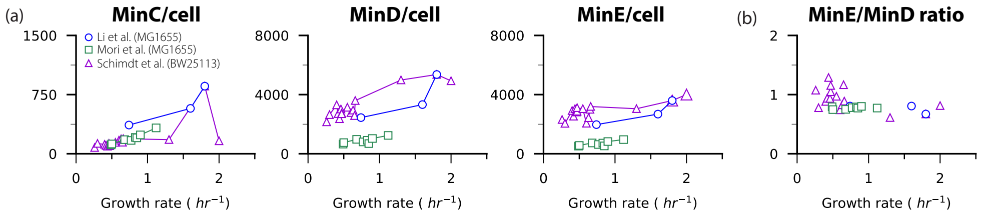

Figure S2. Min protein copy number per cell and MinE/MinD ratio under different growth rates. (a) MinC, MinD and MinE protein copy number per cell under different growth rates. (b) MinE/MinD ratio calculated from panel (a)

not measure the Min proteins. Ribosome profiling is generally more reliable, especially for small copy-number proteins. Mass-spec data are relatively reliable if the protein copy numbers are $10^3$ or more.

We compiled the total copy numbers of MinC, MinD, and MinE per cell vs. growth rate data from the three papers [see Fig. S2a]. From the protein copy number data, we calculated the number ratio between MinE and MinD. Although the total amount of Min proteins increases as the cell grows faster, the MinE/MinD ratio remains surprisingly constant around value 1 [see Fig. S2b].

#### 2.2 Min protein concentration

Next, we calculated the Min protein concentrations for each data set by normalizing the protein copy number using inferred cell sizes [ref. Cell size paper] at different growth rates Ref. [5]. To better compare the Min concentrations between different sources, we chose dataset acquired from the same E.coli background (MG1655). However, due to different protein quantification methods, the Min protein concentrations differ greatly between the two published MG1655 datasets. Therefore, to compare the fold-change and the trend of the Min protein concentration variation between datasets, we first normalized the data point from Mori et al. near growth rate = 0.74/hr by the data point from Li et al. around the same growth rate, and then use that same normalization factor to correct for the rest of the data points from Mori et al. Using the same normalization method, we overlaid our fluorescence measurements to the published data set [see Fig S3e-f].

For both E.coli strain backgrounds, the MinD and MinE protein concentrations decrease as the growth rate increases [see Fig S3b-c], which is expected for constitutive promoters Ref. [6]. The decreasing trend is less clear for MinC concentrations, as only the data set from Li et al. exhibits an obvious decrease in MinC [see Fig S3a,d]. The fluorescence measurements from this study overlay very well with the other two MG1655 datasets, with a similar trend and fold change as the cell transition from slow growth to fast growth environments.

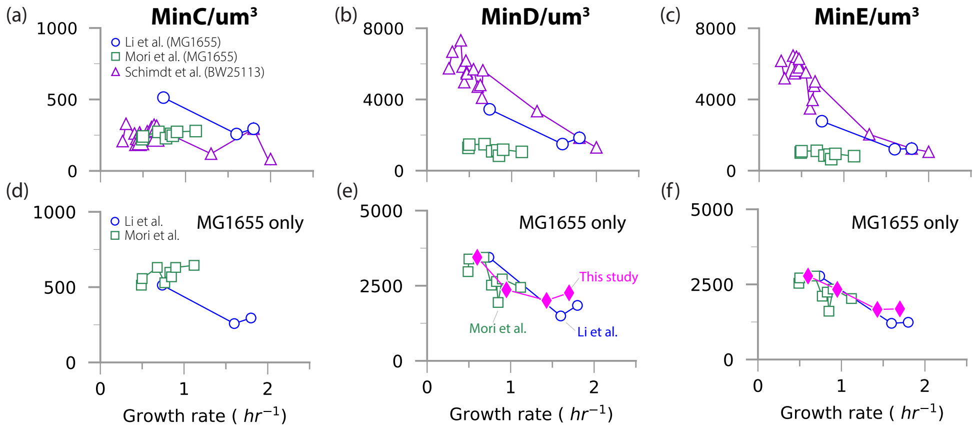

Figure S3. Absolute Min protein concentrations for different growth rates. (a) MinC protein concentrations calculated from Fig. S2. (b) MinD protein concentrations calculated from Fig. S2. (c) MinE protein concentrations calculated from Fig. S2. (d) Normalized MinC protein concentrations for MG1655 background only. (e) Normalized MinD protein concentrations overlaid with data from this study. (f) Normalized MinE protein concentrations overlaid with data from this study. The experiment data points are corrected for the mCherry maturation.

### 3 Min protein fluorescent measurement maturation correction

#### 3.1 GFP-MinD and MinE mCherry maturation curves

In E.coli cells, It has been shown that the two widely used fluorescent markers GFP and mCherry show highly variable maturation times under different growth rates, and mCherry exhibits a much longer maturation time compared to GFP in general [7]. Thus, the direct fluorescent readout does not reflect the actual amount of proteins within the cell and needs to be corrected to take into account the Min proteins associated with unmatured fluorescent markers.

We examined the maturation of the sfGFP Ref. [8] and mCherry Ref. [9] used in this study under four different growth conditions, including the fast and slow growth conditions presented on the experimental phase diagram. To maintain a steady 37C temperature during fluorescence maturation measurement, cell cultures taken from the 37C water bath were immediately transferred and prepared in a temperature-controlled environmental room (Darwin Chambers Company), within which all fluorescence imaging equipment is contained.

In Fig S4, we show three types of time traces. GFP from GFP-MinD (translational reporter), mCherry from MinE mCherry (transcriptional reporter), and their ratio. Cell growth was arrested at time 0 minutes by the addition of chloramphenicol. The total GFP-MinD signal changes negligibly, indicating the fast maturation of sfGFP. By contrast, the mCherry signal increases steadily, reaching a plateau.

##### 3.2 Two-step correction for Min protein measurement

Notice the fold changes are growth-rate dependent because a larger fraction of fluorescent proteins should be in a pre-maturation state in fast-growing cells. Indeed, for the doubling time of 25 minutes, the increase in MinE mCherry signal is 1.72 fold, whereas, at the doubling time of 55 minutes, the increase is 1.33 fold. Another important feature emerges from this carefully controlled protein maturation experiment. The total integrated signal for GFP-MinD and MinE mCherry increases as the growth rate increases. They increase by a similar fold, consistent with the published data in Figure 2c.

Using the results in Fig S4, we plot the MinD/MinE ratio vs. growth rate before and after maturation time correction [Fig S5a]. The most encouraging feature here is that the MinD/MinE is virtually flat, as expected for two genes controlled under the same operon. Furthermore, since the MinD/MinE values from the three independent studies are consistent with each other to be MinD/MinE $\approx 1$, we can constrain our measured MinD/MinE (in arbitrary unit) to be MinD/MinE $= 1$ [Fig S5b].

##### 3.3 MinE-MinD phase diagram with fluorescence maturation correction

Knowing how much to correct for the specific growth conditions used for the old phase diagram (MOPS glycerol minimum and MOPS rich glycerol), we can correct the mCherry value in each growth condition for every point on the previous phase diagram. For example, according to Figure S4 and S5, we must rescale the mCherry signal by 1.72 and 1.33 fold in fast and slow growth conditions, respectively. As for the wild-type expression levels, we measure them directly, as shown in Figure S4.

Since Fig S2b shows that the MinD/MinE ratio remains roughly around value 1 for different growth conditions, we also did a second step correction: rescaling the GFP-MinD value for all the points by the same factor such that the wild-type cell at slow growth condition (first data point in Fig S4b after correction) lies precisely on the diagonal line. By doing that, the x and y axes now share the same arbitrary unit.

#### 4 Cephalexin treatment: decouple cell length from the Min protein levels

The *in vivo* Min protein dynamic patterns crucially depend on the cell length. Intuitively, E.coli cells that are shorter than the intrinsic wavelength of the patterns are unlikely to exhibit traveling waves or standing waves due to the reflection of the wavefront at cell poles before it can travel further. Thus, the experimental phase diagram Figure 2e contains information about the dynamic pattern and the cell morphology as the Min protein concentration varies. For instance, the traveling wave and standing

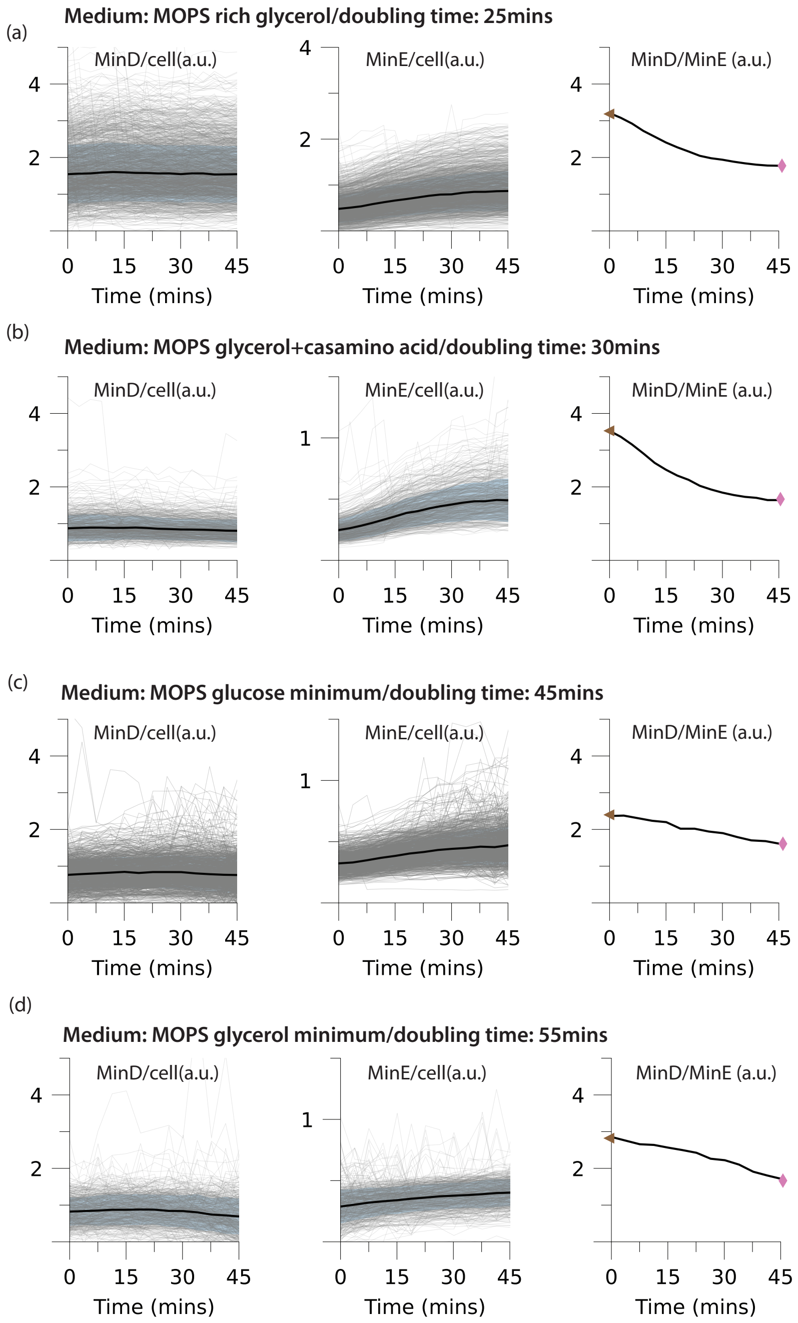

Figure S4. Maturation curves for GFP-MinD and MinE mCherry (wild-type strain SJ1695) at different growth conditions. Cells were treated with chloramphenicol 2 mins prior to t=0 mins shown on the plots. (a) MOPS Rich Glycerol. (b) MOPS glycerol+casamino acid. (c) MOPS glucose minimal. (d) MOPS glycerol minimal.

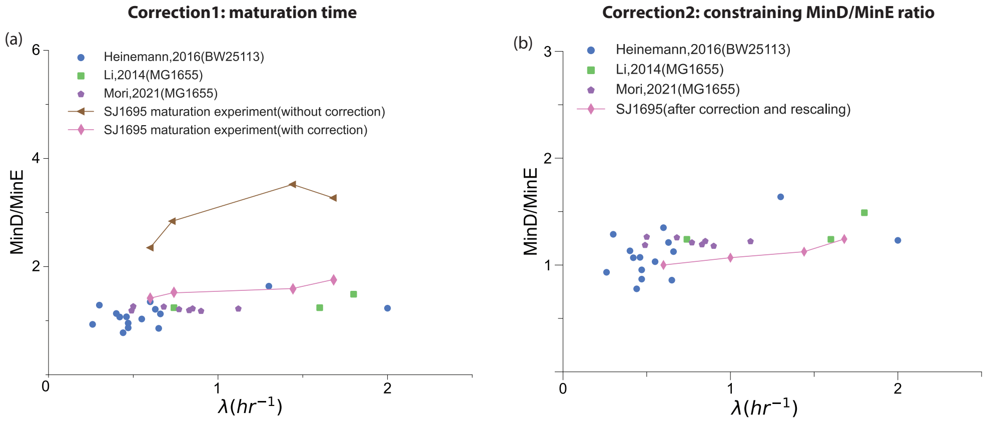

Figure S5. two-step correction procedure used for phase diagram that takes into account fluorophore maturation. (a) Correcting MinD/MinE for maturation time. Brown and pink data points correspond to the time points marked in Fig S4 of the MinD/MinE measurement. (b) We further constrain MinD/MinE = 1 at MOPS glycerol minimum growth condition (doubling time 55mins).

wave regimes are predominately observed in more filamentous cells, whereas the oscillation patterns are mostly found in normal-sized cells. However, the cell morphology becomes a confounding factor when comparing the experiment and the theoretical model because the cell length is coupled to the *in vivo* Min protein concentrations and thus the dynamic patterns.

We decoupled the *in vivo* pattern dependence on the cell length using a sub-lethal dosage of cephalexin (5ng/ml) to inhibit cell division, inducing cell filamentation for conditions that originally produce shorter cells (L<20um) in oscillation and standing wave regimes [Fig S7]. For each condition, when cell length is increased at constant Min protein expression levels, the *in vivo* pattern can become either standing or traveling wave [Fig S7a, SI movies S6,S7]. We first induce filamentation for cells within the oscillation regime and found that once the cell length is decoupled from the Min protein expression levels, the majority of the oscillation pattern in normal cells became standing wave patterns in filamentous cells[Fig S7b,c]. In Fig S7c we showed the representative kymographs for cells before cephalexin treatment and several cell doubling times after the treatment. Interestingly, the oscillation patterns at high MinD and MinE concentrations become traveling wave patterns when cells become filamentous, indicating a downward extension of the upper right traveling-wave regime. We also treat cells in standing wave regimes with cephalexin. When MinD expression levels are low, the standing wave pattern in un-induced conditions remains standing wave even though the cell lengths are considerably increased[SI movie S9]. However, the pattern becomes traveling waves when the MinD expression level is higher [SI movie S9].

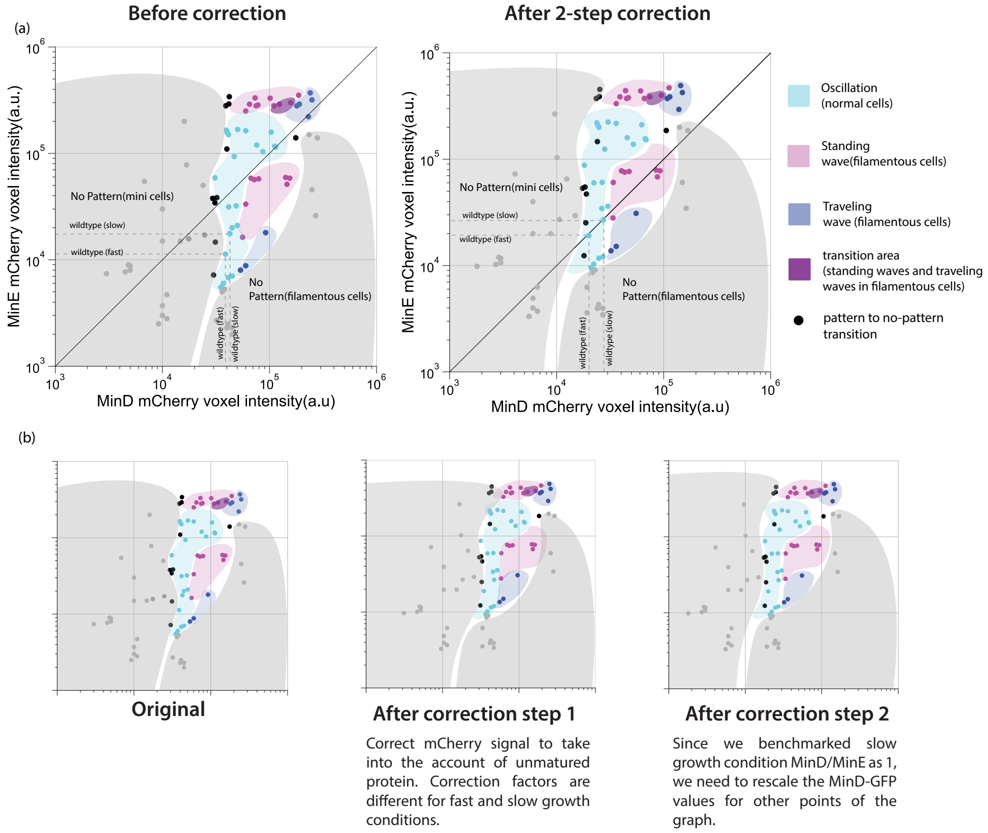

Figure S6. Correcting [MinE] vs. [MinD] phase diagram for fluorophore maturation. (a) phase diagram before and after the two-step correction. (b) step-by-step correction following the same procedure as in Fig S5

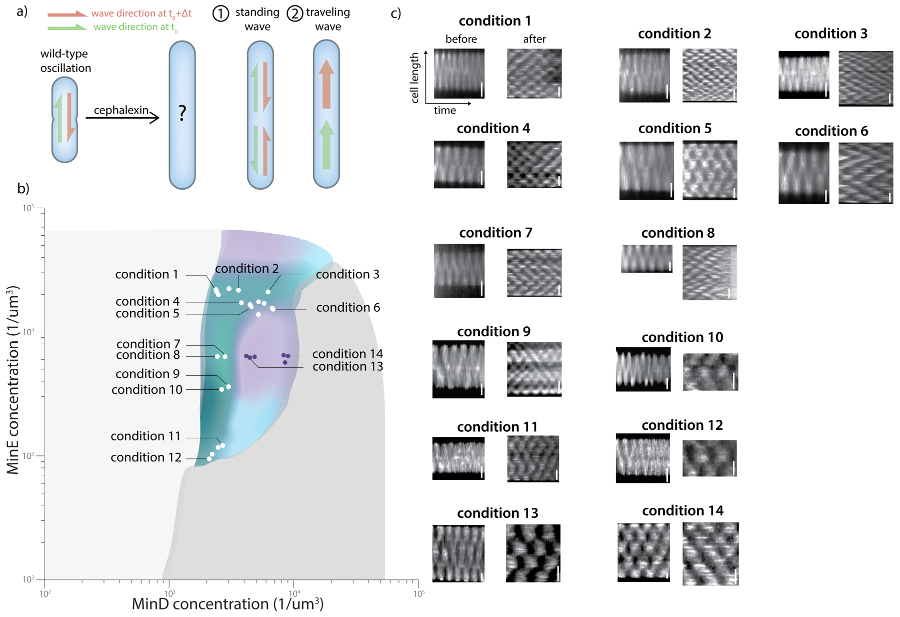

Figure S7. Decouple cell length constraint on observed *in vivo* patterns. (a) Oscillation pattern in normal-sized cells can become traveling or standing waves depending on the Min protein concentrations. (b) Conditions at which the cell filamentation is induced. (c) representative kymographs at each condition before and after cephalexin treatment (scale bar for before-treatment is 1um; scale bar for after-treatment is 10um except for conditions 10/12 which have scale bar of 5um).

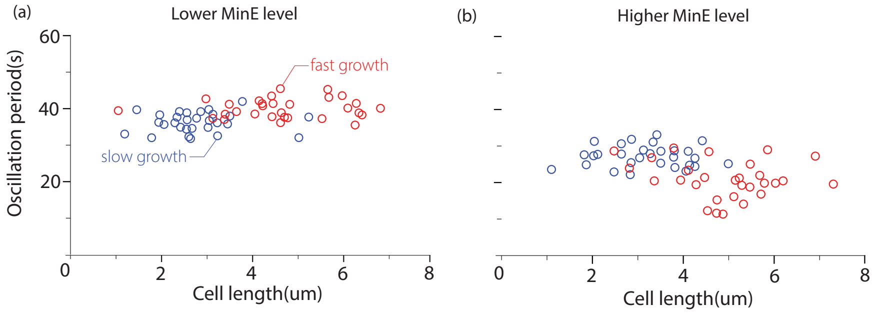

Figure S8. Oscillation period for fast and slow growth conditions at similar Min protein levels (a)Period measurement of fast- and slow-growing cells for condition 9/10 in Fig S7b and wild-type expression levels, each data point is a single cell (b)Period measurement of fast- and slow-growing cells for condition 2/4/5 in Fig S7b, each data point is a single cell

### 5 The Min system shows similar oscillation period at similar Min protein expression levels in different growth conditions

In Fig.2e, we did not distinguish the fast and slow growth conditions in the pattern formation phase diagram and assumed the pattern formation regime remains the same for both fast and slow growth conditions. The two most prominent features of the Min system are its wavelength and period. If both features are only affected by the *in vivo* Min protein levels rather than by the growth conditions, then we can merge the two different growth conditions in the phase diagram.

Besides the pattern wavelength measurement (Fig.4c) which shows a striking constancy across different Min protein expression levels, we also measured the period of oscillation in fast and slow growth conditions at similar Min protein expression levels. Although the fast-growing cells are longer overall than the slow-growing cells, their oscillation periods are fairly similar at similar Min protein levels [Fig. S8]. Notice that at higher MinE expression levels, oscillations in both fast and slow growth conditions show decreased periods [Fig. S8] compared to those at lower MinE expression levels. This observation is also consistent with the model prediction in Fig.4c.

### 6 Comparison between the experiment and theoretically predicted pattern formation regime

When comparing the experiment and theory phase diagrams, it is obvious that the theoretical pattern formation regime extends much more toward high MinD levels than the experimental one (Fig.4b). As observed in the experiment, when MinD levels are

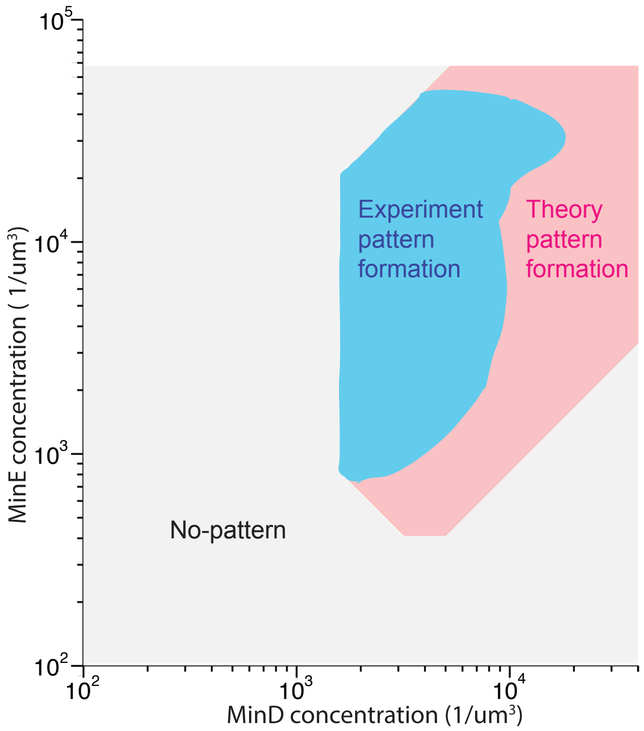

Figure S9. Comparison between experiment and theory pattern formation regimes. The theoretically predicted pattern formation regime extends beyond the right boundary of the experimentally obtained one. The Min system transitions into predominantly no-pattern when MinD levels exceed 20000 per cubic micron.

tuned above 20000 per cubic micron, the traveling wave pattern transitions into the no-pattern regime. In contrast, the theory predicts the existence of the traveling wave pattern above that MinD level[Fig. S9].

We attribute this discrepancy to the experimentally observed MinD aggregation at high MinD levels, which is not considered in the MinE-switch model. At high MinD levels, the localized GFP-MinD signal is not as evenly spread as the signal at lower MinD levels, and GFP-MinD usually forms small bright aggregations as they move along the cell (SI movie2). This aggregation effect of GFP-MinD is more prominent in the no-pattern regime at high MinD levels (SI movie5).

### 7 Models for the *E. coli* Min system in cellular geometry

The basic interaction motives between MinD and MinE have been employed to explain Min protein patterns by a reaction–diffusion mechanism coupling their attachment onto and detachment from the cell membrane to cytosolic and membrane diffusion [10]. Several models that focus on different aspects of the reaction network of the Min proteins have been proposed. The mathematical formulation of the minimal model without the MinE switch ("skeleton model") [11, 12] and with the MinE switch ("switch model") [13] used to explain the experimental data obtained in this study is given in Materials and Methods. We restate the reaction–diffusion equations here in a general form which we use below to reduce the dynamics to a one-dimensional description. We also give the expressions for the general terms in the skeleton and switch models.

To simulate the formation of protein patterns in reaction–diffusion models, one describes the different protein states by their spatiotemporal concentration within the cytosol and on the cell membrane [14]. We model the cellular geometry by a spherocylinder consisting of a cylinder of length $L$ and radius $R$ and two spherical caps at both ends with the same radius $R$ [see Fig. S10(a), $2+3$D geometry]. The cell cytosol corresponds to the bulk of the spherocylinder. The cell membrane is modeled as the surface of the spherocylinder.

The reaction–diffusion dynamics of the proteins follows a set of partial differential equations for all the protein concentrations. The protein concentrations $\mathbf{c}(\mathbf{x}, t)$ in the cytosol undergo diffusion in the 3D cell interior and fulfill

$$\partial_t \mathbf{c} = \mathbf{D}_\mathrm{c} \nabla^2 \mathbf{c} + \mathbf{r}_\mathrm{c}(\mathbf{c}). \tag{1}$$

The cytosolic diffusion coefficients of the different protein species are contained in the diagonal matrix $\mathbf{D}_\mathrm{c}$. We neglect any inhomogeneities in the cytosol, e.g., due to the nucleoid, and approximate the protein motion by average, spatially uniform diffusion constants. The reaction term $\mathbf{r}_\mathrm{c}(\mathbf{c})$ describes the conversion reactions between the cytosolic species. The membrane concentrations $\mathbf{m}(\mathbf{x}, t)$ are similarly determined by

$$\partial_t \mathbf{m} = \mathbf{D}_\mathrm{m} \nabla^2_\mathrm{m} \mathbf{m} + \mathbf{r}_\mathrm{m}(\mathbf{m}, \mathbf{c}|_\mathrm{m}). \tag{2}$$

Here, the diagonal matrix $\mathbf{D}_\mathrm{m}$ contains the membrane diffusion constants, and $\nabla^2_\mathrm{m}$ denotes the Laplace-Beltrami operator necessary to describe the protein diffusion along the curved cell membrane. The reaction term $\mathbf{r}_\mathrm{m}(\mathbf{m}, \mathbf{c}|_\mathrm{m})$ depends both on the membrane concentrations as well as the cytosolic concentrations at the membrane $\mathbf{c}|_\mathrm{m}$ because it describes attachment onto and detachment from the membrane as well as conversion between membrane species.

The attachment and detachment processes induce protein fluxes in the cytosol normal to the cell membrane [see Fig. S10(d)]. This coupling of the cytosolic concentrations to the membrane dynamics is described by the reactive boundary conditions

$$\mathbf{n} \cdot \mathbf{D}_\mathrm{c} \nabla \mathbf{c}|_\mathrm{m} = \boldsymbol{f}(\mathbf{m}, \mathbf{c}|_\mathrm{m}). \tag{3}$$

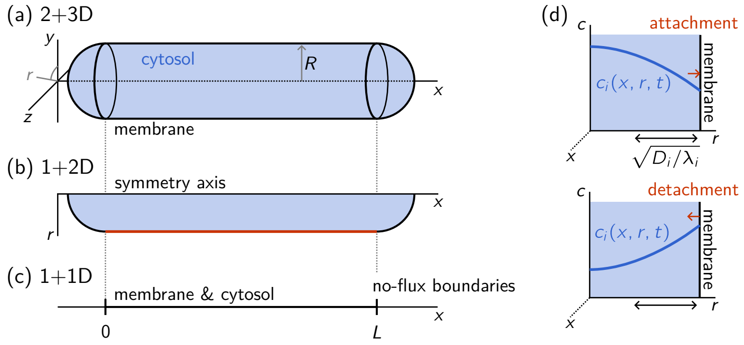

Figure S10. Approximation of the cellular geometry in the reaction–diffusion models. (a) The cell geometry is modeled by a spherocylinder of length $L$ and radius $R$. (b) Employing the rotational symmetry of the analyzed protein patterns, the dynamics can be simulated in a radial slice of the full geometry. To classify the pattern type, the surface concentrations on the membrane are recorded along the red line. (c) If the radius $R$ is small compared to the length scale of the cytosolic gradients perpendicular to the membrane, the reaction–diffusion dynamics may be projected onto a line. The cell poles are represented by no-flux boundary conditions. (d) Illustration of bulk-boundary coupling: Attachment and detachment of proteins onto and from the membrane induce gradients in the cytosolic concentration perpendicular to the membrane. The cytosolic diffusion constant $D_i$ together with the reactive conversion rate in the cytosol $\lambda_i$, i.e., the corresponding reaction rate in $\boldsymbol{r}_c$, sets the typical length scale $\sqrt{D_i/\lambda_i}$ on which the cytosolic concentration $c_i$ varies.

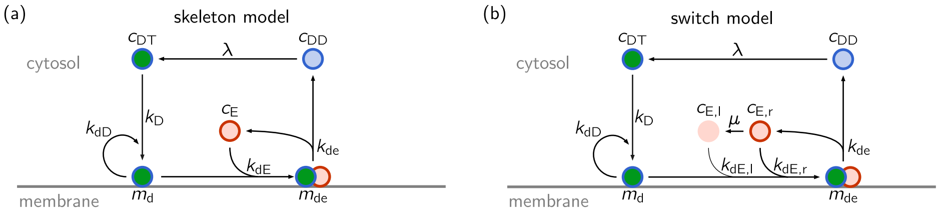

Figure S11. Reaction networks underlying the skeleton (a) and switch (b) models.

The vector $\mathbf{n}$ is the outward-pointing normal vector of the cell membrane. The attachment and detachment flows affecting the cytosolic concentrations $\mathbf{c}$ are contained in the reaction term $\boldsymbol{f}$.

This dynamics conserves the total number of proteins of each species because the proteins only diffuse in the cell and switch between cytosolic and membrane-bound states. For each protein species $i$ we can define the "stoichiometric" vectors $\mathbf{s}_i^{\mathrm{c}}$, $\mathbf{s}_i^{\mathrm{m}}$ such that $\mathbf{s}_i^{\mathrm{c}} \cdot \mathbf{c}$ is the sum of the cytosolic components of species $i$ and $\mathbf{s}_i^{\mathrm{m}} \cdot \mathbf{m}$ is the sum of the membrane components of this species. It follows that the conserved average total protein concentration $\bar{\rho}_i$ of the protein species $i$ is given by

$$\bar{\rho}_i = \frac{1}{V}\left[\int_{\text{cytosol}} \mathrm{d}^3x\, \mathbf{s}_i^{\mathrm{c}} \cdot \mathbf{c}(\mathbf{x},t) + \int_{\text{membrane}} \mathrm{d}^2x\, \mathbf{s}_i^{\mathrm{m}} \cdot \mathbf{m}(\mathbf{x},t)\right], \tag{4}$$

where $V$ is the cytosolic volume, i.e., the volume of the spherocylinder. We now specify the concentration fields and reaction terms for the skeleton and switch models.

#### 7.1 The skeleton model

The skeleton model was introduced in Refs. [11, 12]. This model describes the cytosolic concentrations $\mathbf{c} = (c_{\mathrm{DD}}, c_{\mathrm{DT}}, c_{\mathrm{E}})$ of the ATP- and ADP-bound forms of MinD and cytosolic MinE [see Fig. S11(a)]. In the cytosol, nucleotide exchange reactivates ADP-bound MinD. This results in the reaction term

$$\mathbf{r}_{\mathrm{c}}(\mathbf{c}) = \begin{pmatrix} -\lambda c_{\mathrm{DD}} \\ \lambda c_{\mathrm{DD}} \\ 0 \end{pmatrix}. \tag{5}$$

On the membrane, the concentrations of MinD and MinDE complexes $\mathbf{m} = (m_{\mathrm{d}}, m_{\mathrm{de}})$ are modeled which undergo the reactions

$$\mathbf{r}_{\mathrm{m}}(\mathbf{m},\mathbf{c}) = \begin{pmatrix} (k_{\mathrm{D}} + k_{\mathrm{dD}} m_{\mathrm{d}}) c_{\mathrm{DT}} - k_{\mathrm{dE}} c_{\mathrm{E}} m_{\mathrm{d}} \\ k_{\mathrm{dE}} c_{\mathrm{E}} m_{\mathrm{d}} - k_{\mathrm{de}} m_{\mathrm{de}} \end{pmatrix}. \tag{6}$$

The bulk-boundary coupling occurs via the reaction term

$$\boldsymbol{f}(\mathbf{m}, \mathbf{c}) = \begin{pmatrix} k_{\mathrm{de}} m_{\mathrm{de}} \\ -(k_{\mathrm{D}} + k_{\mathrm{dD}} m_{\mathrm{d}}) c_{\mathrm{DT}} \\ -k_{\mathrm{dE}} m_{\mathrm{d}} c_{\mathrm{E}} + k_{\mathrm{de}} m_{\mathrm{de}} \end{pmatrix} . \tag{7}$$

Moreover, the diffusion matrices are chosen as

$$\mathbf{D}_{\mathrm{c}} = \mathrm{diag}(D_{\mathrm{D}}, D_{\mathrm{D}}, D_{\mathrm{E}}), \qquad \mathbf{D}_{\mathrm{m}} = \mathrm{diag}(D_{\mathrm{d}}, D_{\mathrm{de}}). \tag{8}$$

Finally, the stoichiometric vectors are

$$\mathbf{s}_{\mathrm{D}}^{\mathrm{c}} = \begin{pmatrix} 1 \\ 1 \\ 0 \end{pmatrix}, \qquad \mathbf{s}_{\mathrm{D}}^{\mathrm{m}} = \begin{pmatrix} 1 \\ 1 \end{pmatrix}, \qquad \mathbf{s}_{\mathrm{E}}^{\mathrm{c}} = \begin{pmatrix} 0 \\ 0 \\ 1 \end{pmatrix}, \qquad \mathbf{s}_{\mathrm{E}}^{\mathrm{m}} = \begin{pmatrix} 0 \\ 1 \end{pmatrix}. \tag{9}$$

This model conserves the average total MinD and MinE concentrations $\bar{\rho}_{\mathrm{D}}$ and $\bar{\rho}_{\mathrm{D}}$:

$$\bar{\rho}_{\mathrm{D}} = \frac{1}{V} \left[ \int_{\mathrm{cytosol}} \mathrm{d}^3 x \, \big( c_{\mathrm{DD}}(\mathbf{x}, t) + c_{\mathrm{DT}}(\mathbf{x}, t) \big) + \int_{\mathrm{membrane}} \mathrm{d}^2 x \, \big( m_{\mathrm{d}}(\mathbf{x}, t) + m_{\mathrm{de}}(\mathbf{x}, t) \big) \right], \tag{10}$$

$$\bar{\rho}_{\mathrm{E}} = \frac{1}{V} \left[ \int_{\mathrm{cytosol}} \mathrm{d}^3 x \, c_{\mathrm{E}}(\mathbf{x}, t) + \int_{\mathrm{membrane}} \mathrm{d}^2 x \, m_{\mathrm{de}}(\mathbf{x}, t) \right]. \tag{11}$$

###### 7.1.1 Parameter choice

As stated in Material and Methods, we use the skeleton model with the parameters determined in Ref. [12]. However, the nonlinear rates are rescaled to observe pattern formation at comparable values with the phase diagram found experimentally in this article (see Fig. 4 a). For the reader's convenience, we summarize the parameter values in Table 1.

##### 7.2 The switch model

A striking feature of the skeleton model is that it only forms patterns if the average total concentration of MinE is lower than or roughly equal to the average concentration of MinD proteins (see Fig. 4 a in the main text and Ref. [12, 13]). In contrast, the experiments of this study show that patterns form over a range of more than one order of magnitude of MinE concentrations.

A similar robustness of the Min protein patterns was observed in *in vitro* experiments in Ref. [13] The authors showed that the robustness is due to the conformational switch of MinE dimers between a latent (closed) and reactive (open) state. To describe this conformational switch, the "switch model" was introduced as an extension of the skeleton model [13] [see Fig. S11(b)]. Adding a latent MinE conformation to the skeleton model allows this inert state to act as a buffer at high MinE concentrations, which effectively reduces the active MinE concentration and confers robustness on the Min protein patterns.

| Parameter | Value | Unit |
|---|---|---|
| $D_\mathrm{D}$ | 16 | $\mu\mathrm{m}^2\mathrm{s}^{-1}$ |
| $D_\mathrm{E}$ | 10 | $\mu\mathrm{m}^2\mathrm{s}^{-1}$ |
| $D_\mathrm{d}$ | 0.013 | $\mu\mathrm{m}^2\mathrm{s}^{-1}$ |
| $D_\mathrm{de}$ | 0.013 | $\mu\mathrm{m}^2\mathrm{s}^{-1}$ |
| $k_\mathrm{D}$ | 0.1 | $\mu\mathrm{m}\,\mathrm{s}^{-1}$ |
| $k_\mathrm{dD}$ | 0.108/60 | $\mu\mathrm{m}^3\mathrm{s}^{-1}$ |
| $k_\mathrm{dE}$ | 0.435/60 | $\mu\mathrm{m}^3\mathrm{s}^{-1}$ |
| $k_\mathrm{de}$ | 1.925 | $\mathrm{s}^{-1}$ |
| $\lambda$ | 6 | $\mathrm{s}^{-1}$ |
| $\bar{\rho}_\mathrm{D}$ | varied | $\mu\mathrm{m}^{-3}$ |
| $\bar{\rho}_\mathrm{E}$ | varied | $\mu\mathrm{m}^{-3}$ |

Table 1. Scaled parameters based on Ref. [12] used here for the skeleton model [see Fig. S11(a)]. Because the experiments presented in this article were performed at 37 °C, we evaluated the hydrolysis rate $k_\mathrm{de}$ at this temperature.

While the switch model describes the same protein concentrations on the membrane as the skeleton model, two distinct concentration fields $c_\mathrm{E,r}$ and $c_\mathrm{E,l}$ are introduced in the cytosol for the reactive and latent MinE conformations. This results in $\mathbf{c} = (c_\mathrm{DD}, c_\mathrm{DT}, c_\mathrm{E,r}, c_\mathrm{E,l})$. The cytosolic reaction term is modified as

$$\mathbf{r}_\mathrm{c}(\mathbf{c}) = \begin{pmatrix} -\lambda c_\mathrm{DD} \\ \lambda c_\mathrm{DD} \\ -\mu c_\mathrm{E,r} \\ \mu c_\mathrm{E,r} \end{pmatrix}, \tag{12}$$

where $\mu$ describes the rate of the conformational switch. The membrane reactions in the switch model read

$$\mathbf{r}_\mathrm{m}(\mathbf{m}, \mathbf{c}) = \begin{pmatrix} (k_\mathrm{D} + k_\mathrm{dD} m_\mathrm{d}) c_\mathrm{DT} - (k_\mathrm{dE,r} c_\mathrm{E,r} + k_\mathrm{dE,l} c_\mathrm{E,l}) m_\mathrm{d} \\ (k_\mathrm{dE,r} c_\mathrm{E,r} + k_\mathrm{dE,l} c_\mathrm{E,l}) m_\mathrm{d} - k_\mathrm{de} m_\mathrm{de} \end{pmatrix}. \tag{13}$$

The attachment-detachment flows are given by

$$\boldsymbol{f}(\mathbf{m}, \mathbf{c}) = \begin{pmatrix} k_\mathrm{de} m_\mathrm{de} \\ -(k_\mathrm{D} + k_\mathrm{dD} m_\mathrm{d}) c_\mathrm{DT} \\ -k_\mathrm{dE,r} c_\mathrm{E,r} m_\mathrm{d} + k_\mathrm{de} m_\mathrm{de} \\ -k_\mathrm{dE,l} c_\mathrm{E,l} m_\mathrm{d} \end{pmatrix}. \tag{14}$$

The matrix of cytosolic diffusion constants is extended to

$$\mathbf{D}_\mathrm{c} = \mathrm{diag}(D_\mathrm{D}, D_\mathrm{D}, D_\mathrm{E}, D_\mathrm{E}). \tag{15}$$

Moreover, the stoichiometric vectors read

$$\mathbf{s}_\mathrm{D}^\mathrm{c} = \begin{pmatrix} 1 \\ 1 \\ 0 \\ 0 \end{pmatrix}, \qquad \mathbf{s}_\mathrm{D}^\mathrm{m} = \begin{pmatrix} 1 \\ 1 \end{pmatrix}, \qquad \mathbf{s}_\mathrm{E}^\mathrm{c} = \begin{pmatrix} 0 \\ 0 \\ 1 \\ 1 \end{pmatrix}, \qquad \mathbf{s}_\mathrm{E}^\mathrm{m} = \begin{pmatrix} 0 \\ 1 \end{pmatrix}. \tag{16}$$

| Parameter | Value | Unit |
|---|---|---|
| $D_{\mathrm{D}}$ | 16 | $\mu\mathrm{m}^2\mathrm{s}^{-1}$ |
| $D_{\mathrm{E}}$ | 10 | $\mu\mathrm{m}^2\mathrm{s}^{-1}$ |
| $D_{\mathrm{d}}$ | 0.05 | $\mu\mathrm{m}^2\mathrm{s}^{-1}$ |
| $D_{\mathrm{de}}$ | 0.05 | $\mu\mathrm{m}^2\mathrm{s}^{-1}$ |
| $k_{\mathrm{D}}$ | 0.3 | $\mu\mathrm{m}\,\mathrm{s}^{-1}$ |
| $k_{\mathrm{dD}}$ | $7.5{\cdot}10^{-4}$ | $\mu\mathrm{m}^3\mathrm{s}^{-1}$ |
| $k_{\mathrm{dE,r}}$ | 0.75 | $\mu\mathrm{m}^3\mathrm{s}^{-1}$ |
| $k_{\mathrm{dE,l}}$ | $5{\cdot}10^{-6}$ | $\mu\mathrm{m}^3\mathrm{s}^{-1}$ |
| $k_{\mathrm{de}}$ | 1 | $\mathrm{s}^{-1}$ |
| $\lambda$ | 5 | $\mathrm{s}^{-1}$ |
| $\mu$ | 20 | $\mathrm{s}^{-1}$ |
| $\bar{\rho}_{\mathrm{D}}$ | varied | $\mu\mathrm{m}^{-3}$ |
| $\bar{\rho}_{\mathrm{E}}$ | varied | $\mu\mathrm{m}^{-3}$ |
| $R$ | 0.5 | $\mu\mathrm{m}$ |
| $L$ | 50 | $\mu\mathrm{m}$ |

Table 2. Parameters used here for the switch model [see Fig. S11(b)] to describe the experimental concentration phase diagram.

In the switch model, the conservation of the average total MinD and MinE concentrations reads

$$\bar{\rho}_{\mathrm{D}} = \frac{1}{V}\left[\int_{\mathrm{cytosol}} \mathrm{d}^3x \left(c_{\mathrm{DD}}(\mathbf{x},t) + c_{\mathrm{DT}}(\mathbf{x},t)\right) + \int_{\mathrm{membrane}} \mathrm{d}^2x \left(m_{\mathrm{d}}(\mathbf{x},t) + m_{\mathrm{de}}(\mathbf{x},t)\right)\right], \tag{17}$$

$$\bar{\rho}_{\mathrm{E}} = \frac{1}{V}\left[\int_{\mathrm{cytosol}} \mathrm{d}^3x \left(c_{\mathrm{E,r}}(\mathbf{x},t) + c_{\mathrm{E,l}}(\mathbf{x},t)\right) + \int_{\mathrm{membrane}} \mathrm{d}^2x \, m_{\mathrm{de}}(\mathbf{x},t)\right]. \tag{18}$$

##### 7.2.1 Parameter choice

The reaction parameters employed in the switch model are discussed in Materials and Methods section. We restate the parameter values in Table 2.

### 8 Dimensional reduction of the cellular geometry

Because the reaction rates in the Min reaction network are not known from experiments, a large parameter study is necessary to identify a parameter region that reproduces the experimentally observed pattern types. For each parameter set the phase diagram of the average total MinD and MinE concentrations has to be simulated to analyze the patterns in the different concentration regions. To reduce the computational complexity, we reduce the full cellular geometry by using the cylindrical symmetry of the cell geometry.

#### 8.1 Rotational symmetry

The *E. coli* cell geometry is (approximately) rotationally symmetric around the long axis of the bacteria. This is reflected by the rotational symmetry of the spherocylinder in our reaction–diffusion model. Moreover, the studied protein patterns are rotationally symmetric around the bacterial long axis as well after the initial transient of pattern growth. Thus, the protein concentration fields only depend on the axial position $x$ along the long axis of the cell and on the radial position $r$:

$$\mathbf{c} = \mathbf{c}(x, r, t), \tag{19}$$

$$\mathbf{m} = \mathbf{m}(x, t). \tag{20}$$

Here, the protein concentrations on the membrane $\mathbf{m}$ are given in terms of a Monge parametrization.

The assumption of radial symmetry allows the simulation of the full dynamics including the bulk-boundary coupling between the cytosol and membrane as a 1+2D system instead of simulating the full 2+3D dynamics [see Fig. S10(b)]. Employing the form of the Laplacian in cylindrical coordinates, the cytosolic dynamics reduce to

$$\partial_t \mathbf{c}(x, r, t) = \mathbf{D}_\mathrm{c} \left( \partial_x^2 + \frac{1}{r}\partial_r + \partial_r^2 \right) \mathbf{c} + \mathbf{r}_\mathrm{c}(\mathbf{c}). \tag{21}$$

To determine the corresponding membrane dynamics, the Laplace-Beltrami operator has to be expressed in radial coordinates. The Laplace-Beltrami operator can be expressed using the surface derivative $\nabla_\mathrm{m} = (\mathbf{1} - \mathbf{n}\mathbf{n}^T)\nabla$ where $\mathbf{n}$ is the surface unit normal vector. The matrix $(\mathbf{1} - \mathbf{n}\mathbf{n}^T)$ projects the derivative onto the tangent plane of the surface. The Laplace-Beltrami operator then reads

$$\nabla_m^2 \bullet = (\mathbf{1} - \mathbf{n}\mathbf{n}^T)\nabla \left[ (\mathbf{1} - \mathbf{n}\mathbf{n}^T)\nabla \bullet \right]. \tag{22}$$

Expressing every term in cylindrical coordinates and using that the concentration fields are independent of the angular coordinate, one finds from Eq. (2) the reduced membrane dynamics

$$\partial_t \mathbf{m}(x, t) = \mathbf{D}_\mathrm{m}(\nabla_{\mathrm{2D,m}}^2 + \frac{1}{r}\mathbf{e}_r \cdot \nabla_{\mathrm{2D,m}})\mathbf{m} + \mathbf{r}_\mathrm{m}(\mathbf{m}, \mathbf{c}|_\mathrm{m}). \tag{23}$$

The derivative $\nabla_{\mathrm{2D,m}} = (\mathbf{1} - \mathbf{n}\mathbf{n}^T)(\mathbf{e}_r\partial_r + \mathbf{e}_x\partial_x)$ is the tangential derivative of the one-dimensional membrane line in the two-dimensional radial slice $(x, r)$ of the three-dimensional geometry. The vectors $\mathbf{e}_r$ and $\mathbf{e}_x$ are the unit vectors in the radial and axial direction.

#### 8.2 Reduction to a one-dimensional model

In general, the extension of the cytosol in the radial direction cannot be neglected because membrane attachment and detachment induce gradients in the concentrations of the cytosolic protein species normal to the membrane [see Eq. (3) and Fig. S10(d)].

The reduction onto a 1+2D model exactly reproduces these gradients (under the assumption of cylindrically symmetric patterns). If the gradients are shallow compared to the depth of the cytosol, the dynamics can be approximately reduced onto a line neglecting the extension of the cytosol perpendicular to the membrane [see Fig. S10(c)].

The length scale of the cytosolic gradients can be estimated by the typical diffusion length scales set by the ratio of the cytosolic diffusion coefficients and the cytosolic conversion rates $\lambda$ and $\mu$ [14–16]. For the experimentally measured cytosolic diffusion coefficients and the conversion rates employed in our simulations (see Tables 1, 2), the diffusion length scales for MinD and MinE are $\sqrt{D_\mathrm{D}/\lambda} \approx 1.8\,\mu\mathrm{m}$ and $\sqrt{D_\mathrm{E}/\mu} \approx 0.7\,\mu\mathrm{m}$. Thus, the gradients occur on length scales larger than the typical radius of *E. coli* bacteria of about $0.5\,\mu\mathrm{m}$. Therefore, we approximate the cytosolic concentrations as constant normal to the membrane in realistic cell geometries, and the radial direction can be integrated out. The result is a model in a reduced line geometry where both the membrane and cytosolic concentrations only depend on the coordinate $x$ along the long axis of the bacteria [see Fig. S10(c)].

Neglecting the spherical caps at the ends of the spherocylinder used to approximate the cellular geometry, we can integrate the cytosolic concentration fields over cross sections normal to the long axis $x$, and we arrive at

$$\partial_t 2\pi \int_0^R \mathrm{d}r\, r\mathbf{c}(r,x,t) = 2\pi R\,\partial_t \tilde{\mathbf{c}}(x,t) \tag{24}$$

$$= 2\pi R \boldsymbol{f}\big(\mathbf{m}(x,t), \mathbf{c}(R,x,t)\big) + 2\pi R D_\mathrm{c}\partial_x^2\,\tilde{\mathbf{c}} + 2\pi R \mathbf{r}_\mathrm{c}(\tilde{\mathbf{c}}). \tag{25}$$

Here, we introduced the cross-sectional (surface) concentrations $\tilde{\mathbf{c}}(x,t)$ that describe the protein content within a cross-section of the cylinder projected onto the circumference, i.e., on the membrane:

$$\tilde{\mathbf{c}}(x,t) = \frac{1}{2\pi R}\int_0^R \mathrm{d}r\, 2\pi r\mathbf{c}(r,x,t). \tag{26}$$

The reaction term $\mathbf{r}_\mathrm{c}$ is linear for the models we consider in this study. This allows it to be rewritten in terms of the reduced concentrations $\mathbf{c}'(x,t)$.

Under the assumption that the cytosolic gradients normal to the membrane are small in the considered cellular geometry, Eq. (26) simplifies to

$$\mathbf{c}(R,x,t) \approx \frac{2\pi R}{\pi R^2}\mathbf{c}'(x,t) = \frac{\mathbf{c}'(x,t)}{\zeta}, \tag{27}$$

introducing the bulk-surface ratio $\zeta$. Inserting this approximation in Eq. (25), one finds the reduced one-dimensional cytosolic dynamics

$$\partial_t \mathbf{c}' \approx \mathbf{D}_\mathrm{c}\partial_x^2\,\mathbf{c}' + \boldsymbol{f}\left(\mathbf{m}, \frac{\mathbf{c}'}{\zeta}\right) + \mathbf{r}_\mathrm{c}(\mathbf{c}') = \mathbf{D}_\mathrm{c}\partial_x^2\,\mathbf{c}' + \tilde{\boldsymbol{f}}(\mathbf{m},\mathbf{c}') + \mathbf{r}_\mathrm{c}(\mathbf{c}'). \tag{28}$$

By the same approximation, one finds from Eq. (2) the reduced membrane dynamics

$$\partial_t \mathbf{m} \approx \mathbf{D}_\mathrm{m}\partial_x^2\,\mathbf{m} + \mathbf{r}_\mathrm{m}\left(\mathbf{m}, \frac{\mathbf{c}'}{\zeta}\right) = \mathbf{D}_\mathrm{m}\partial_x^2\,\mathbf{m} + \tilde{\mathbf{r}}_\mathrm{m}(\mathbf{m},\mathbf{c}'). \tag{29}$$

The reaction terms $\tilde{\boldsymbol{f}}$ and $\tilde{\mathbf{r}}_{\mathrm{m}}$ include the bulk-surface ratio in the reaction constants.

For the cylindrical geometry considered the bulk-surface ratio is given by $\zeta = R/2$. In a spherocylinder, the bulk-surface ratio is a little smaller due to the additional membrane area at the cell poles. For short cells of wild-type length $L \approx 3\,\mu\mathrm{m}$ and $R = 0.5\,\mu\mathrm{m}$ this results in $\zeta \approx 0.23\,\mu\mathrm{m}$ instead of $R/2 = 0.25\,\mu\mathrm{m}$. For filamentous cells, the bulk-boundary ratio approaches $R/2$. As the difference is small even for short cells, we use the value $\zeta = R/2 = 0.25\,\mu\mathrm{m}$ throughout this study for all cell lengths. Moreover, the cell poles are not modeled explicitly and only included as no-flux boundary conditions of the one-dimensional domain of length $L$ in the simulation of Eqs. (28), (29).

Finally, in the reduced line dynamics the conservation laws for the average total (surface) concentrations $\bar{\rho}_i$ simply read

$$\bar{\rho}_i = \frac{1}{\zeta L} \int_0^L \mathrm{d}x \left( \mathbf{s}_i^{\mathrm{c}} \cdot \mathbf{c}'(\mathbf{x}, t) + \mathbf{s}_i^{\mathrm{m}} \cdot \mathbf{m}(\mathbf{x}, t) \right). \tag{30}$$

#### 8.3 Numerical simulation

The reduced 1+2D model accounting for the realistic cell geometry is simulated using COMSOL Multiphysics 6.0 & 6.1. The simulations employ a finite-element discretization on a triangular Delaunay mesh with linear Lagrange elements for the cytosol (bulk) and quadratic elements on the membrane (boundary). Initially, the total protein mass is homogeneously distributed in the cytosolic states $c_{\mathrm{DT}}$ and $c_{\mathrm{E}}$ or $c_{\mathrm{E,l}}$ with weak random perturbations (uniformly distributed within $\pm 1\%$ around the homogeneous concentration). The 1+1D dynamics is solved using a uniform finite-differences discretization implemented in Mathematica 13.0 & 13.1 using second-order central differences. In the simulations, we employ the freedom in the choice of units, and the fast-growth wild-type concentrations are normalized to $\bar{\rho}_{\mathrm{D}} = \bar{\rho}_{\mathrm{E}} = 1000\,\mu\mathrm{m}^{-3}$ by scaling the nonlinear rates $k_{\mathrm{dD}}$, $k_{\mathrm{dE}}$, $k_{\mathrm{dE,r}}$, $k_{\mathrm{dE,l}}$ by a factor 1/2. Example notebooks for the simulation, as well as the configuration files for the COMSOL simulation of the switch model, are available at `https://github.com/henrikweyer/Min-in-vivo` [17].

### 9 Linear stability analysis of the homogeneous steady state

Denk et al. [13] used a linear stability analysis to show that the introduction of the MinE switch in the switch model increases the range of concentration ratios between MinD and MinE for which patterns can be observed in *in vitro* geometry.

The linear stability analysis of the homogeneous steady state describes the growth rates for weak perturbations around the homogeneous steady state. In the reduced

1+1D dynamics, the homogeneous steady state $(\mathbf{m}^*, \mathbf{c}'^*)$ is determined by

$$0 = \tilde{\boldsymbol{f}}(\mathbf{m}^*, \mathbf{c}'^*) + \mathbf{r}_\mathrm{c}(\mathbf{c}'^*), \tag{31a}$$
$$0 = \tilde{\mathbf{r}}_\mathrm{m}(\mathbf{m}^*, \mathbf{c}'^*), \tag{31b}$$
$$\rho_i = \mathbf{s}_i^\mathrm{c} \cdot \mathbf{c}'^*(\mathbf{x}, t) + \mathbf{s}_i^\mathrm{m} \cdot \mathbf{m}^*(\mathbf{x}, t), \tag{31c}$$

for each protein species $i$ (MinD and MinE).

The dynamics of weak perturbations $(\delta\mathbf{m}, \delta\mathbf{c})$ around the homogeneous steady state is determined by the linearized dynamics

$$\partial_t \begin{pmatrix} \delta\mathbf{m} \\ \delta\mathbf{c} \end{pmatrix} = \left[ \begin{pmatrix} \mathbf{D}_\mathrm{m} & 0 \\ 0 & \mathbf{D}_\mathrm{c} \end{pmatrix} \partial_x^2 + \mathbf{J}|_{\mathbf{m}^*, \mathbf{c}'^*} \right] \begin{pmatrix} \delta\mathbf{m} \\ \delta\mathbf{c} \end{pmatrix}, \tag{32}$$

where the Jacobian is given by

$$\mathbf{J} = \begin{pmatrix} \partial_\mathbf{m} \tilde{\mathbf{r}}_\mathrm{m} & \partial_{\mathbf{c}'} \tilde{\mathbf{r}}_\mathrm{m} \\ \partial_\mathbf{m} \tilde{\boldsymbol{f}} & \partial_{\mathbf{c}'} (\tilde{\boldsymbol{f}} + \tilde{\mathbf{r}}_\mathrm{c}) \end{pmatrix}. \tag{33}$$

The eigenmodes of the linearized dynamics Eq. (32) are Fourier modes with wavenumber $q$ and growth rate $\sigma(q)$

$$\begin{pmatrix} \delta\mathbf{m}_q \\ \delta\mathbf{c}_q \end{pmatrix} = \begin{pmatrix} \delta\mathbf{m} \\ \delta\mathbf{c} \end{pmatrix} \mathrm{e}^{\mathrm{i}qx + \sigma(q)t}. \tag{34}$$

The collection of growth rates $\sigma(q)$ is called the dispersion relation. It describes the exponential growth rate of a perturbation mode with wavenumber $q$. The growth rate is determined by inserting the ansatz Eq. (34) into the linearized dynamics Eq. (32). This yields the eigenvalue equation

$$\sigma(q) \begin{pmatrix} \delta\mathbf{m}_q \\ \delta\mathbf{c}_q \end{pmatrix} = \left[ - \begin{pmatrix} \mathbf{D}_\mathrm{m} & 0 \\ 0 & \mathbf{D}_\mathrm{c} \end{pmatrix} q^2 + \mathbf{J}|_{\mathbf{m}^*, \mathbf{c}'^*} \right] \begin{pmatrix} \delta\mathbf{m}_q \\ \delta\mathbf{c}_q \end{pmatrix}. \tag{35}$$

Consequently, the dimension of the eigenvalue problem Eq. (35) determines the maximal number of solutions $\sigma(q)$, giving different branches $\sigma_i(q)$ of the dispersion relation. In the following, we denote by the dispersion relation always the eigenvalue $\sigma_i(q)$ for a certain wavenumber $q$ which has the largest real part.

If the real part of the dispersion relation $\Re[\sigma(q)]$ is positive for any $q > 0$, the homogeneous steady state is laterally unstable. Perturbation modes with the corresponding wavenumber grow and the protein concentrations become inhomogeneous as a concentration pattern grows out of the homogeneous steady state.

#### 9.1 Numerical calculation

The linear stability analysis is performed using Mathematica 13.0 & 13.1. First, Eqs. (31) are solved numerically to determine the homogeneous steady state concentrations $\boldsymbol{m}^*$, $\boldsymbol{c}'^*$. If several homogeneous steady states exist, the homogeneous

steady state is selected that is locally stable, i.e., the one which fulfills $\Re[\sigma(0)] = 0$.*
Then, the eigenvalue problem Eq. (35) is solved for discrete wavenumbers $q$. For each wavenumber, the eigenvalue with the largest real part is selected, and their collection for different wavenumbers gives the numerical dispersion relation. The Mathematica notebooks can be found under `https://github.com/henrikweyer/Min-in-vivo` [17].

#### 9.2 Emergence of robust pattern formation due to the MinE switch

Figure S12 (see also Fig. 4 a in the main text) shows that the MinE switch enhances the pattern robustness in the switch model in the reduced 1+1D geometry, i.e., it enlarges the region of pattern formation with $\Re[\sigma(q)] > 0$ for some wavenumber $q$.† If the MinE recruitment rates $k_{\mathrm{dE,r}}$ and $k_{\mathrm{dE,l}}$ are equal, the skeleton model is recovered. This is seen by defining $c_{\mathrm{E}} = c_{\mathrm{E,r}} + c_{\mathrm{E,l}}$ in the reaction–diffusion equations (see Sec. 7). As the recruitment rate $k_{\mathrm{dE,l}}$ of latent MinE is reduced, the homogeneous steady state becomes laterally unstable at increased total MinE concentrations $\bar{\rho}_{\mathrm{E}}$. The increase of the recruitment rate of reactive MinE $k_{\mathrm{dE,r}}$ ensures that pattern formation is possible even at low total MinE and MinD concentrations. Thus, the pattern robustness can be increased in *in vivo* geometry—modeled here as the reduced 1+1D geometry—similarly as *in vitro* (cf. Ref. [13]).

#### 9.3 Onset of instability in the concentration phase diagram

The onset of lateral instability can be super- or subcritical. In the first case, a patterned steady state only exists in the parameter region of lateral instability. The pattern amplitude decreases to zero continuously as the onset of lateral instability is approached by tuning some parameter. In contrast, if the onset is subcritical, patterns can form already before the onset of lateral instability where the homogeneous steady state is linearly stable. In this multistable regime (both patterned and homogeneous steady states exist), a sufficiently large stimulus is necessary to form a pattern.

McRD systems are known to show wide regimes of stimulus-induced pattern formation [19]. Therefore, we analyze the onset of pattern formation in the switch model as well for the chosen parameter set (cf. Table 2). To this end, the nonlinear dynamics of the 1+1D model is simulated using COMSOL Multiphysics [18] for concentration values just inside the region of lateral instability. The simulation is initialized with al proteins in the cytosolic species and sinusoidal spatial perturbations with the wavelength $2\pi/q_{\mathrm{c}}$ given by the critical wavenumber $q_{\mathrm{c}}$ at the onset of pattern formation (see Sec. 9.4). Then a weak, constant degradation term is introduced at $t = 10^4$s either for $c'_{\mathrm{DD}}$ or $c'_{\mathrm{E,l}}$ that slowly reduces the average total MinD or MinE concentration $\bar{\rho}_{\mathrm{D}}$ or

* Each conservation law leads to one zero eigenvalue at $q = 0$.

† The wavenumber $q$ can only take on multiples of $\pi/L$ on a domain with finite length $L$ and no-flux boundaries. The dispersion relation becomes continuous for $L \to \infty$. The resulting stability boundary estimates the stability region of finite systems with a length $L$ that is large compared to the wavelength of the unstable wave modes. We choose $L = 50\,\mu\mathrm{m}$ in the numerical simulations, and the pattern wavelength $\Lambda \approx 10\,\mu\mathrm{m}$ is significantly shorter.

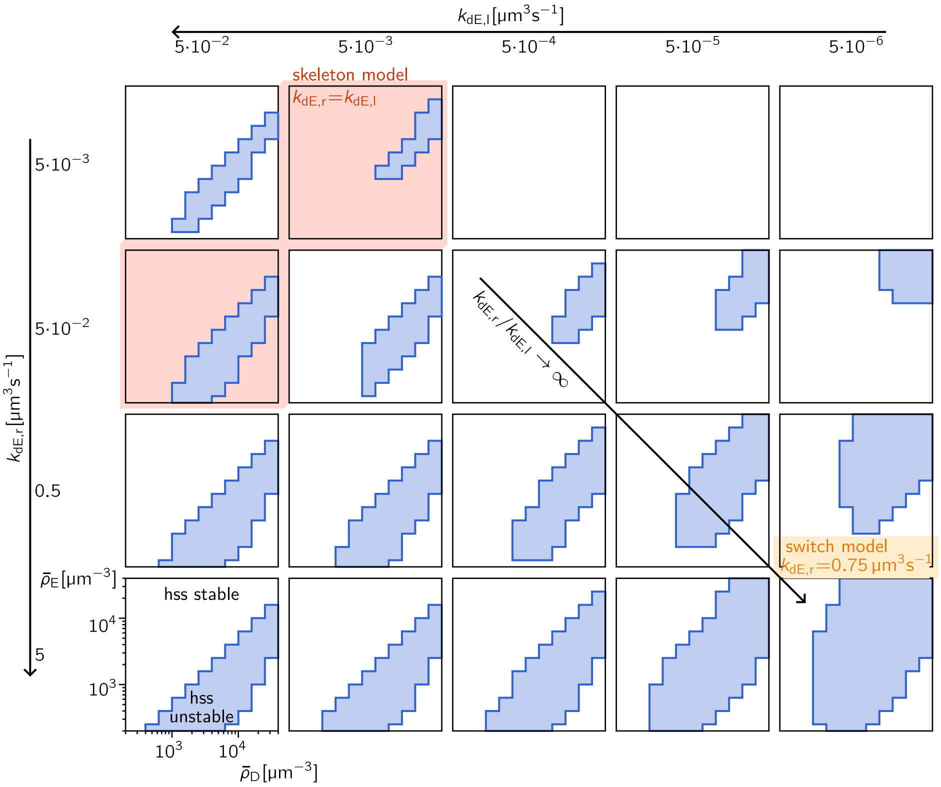

Figure S12. Linear stability analysis of the homogeneous steady state in the switch model. The concentration phase diagrams of the switch model in 1+1D geometry are shown for different combinations of the recruitment rates of the reactive and latent MinE states $k_{\mathrm{dE,r}}$ and $k_{\mathrm{dE,l}}$. The other parameters of the switch model are chosen as given in Table 2. The blue-shaded region denotes the average total MinD and MinE concentrations $\bar\rho_{\mathrm{D}}$ and $\bar\rho_{\mathrm{E}}$ for which the homogeneous steady state is unstable, i.e., $\Re[\sigma(q)] > 0$ for some wavenumber $q$. The same logarithmic concentration scale is used for each phase diagram. The switch model reduces to the skeleton model if $k_{\mathrm{dE,r}} = k_{\mathrm{dE,r}}$ (red-shaded phase diagrams). For comparison with the experimental data, we use $k_{\mathrm{dE,r}} = 0.75\,\mathrm{\mu m^3 s^{-1}}$ and $k_{\mathrm{dE,r}} = 5{\cdot}10^{-6}\mathrm{\mu m^3 s^{-1}}$ (green label; cf. Table 2).

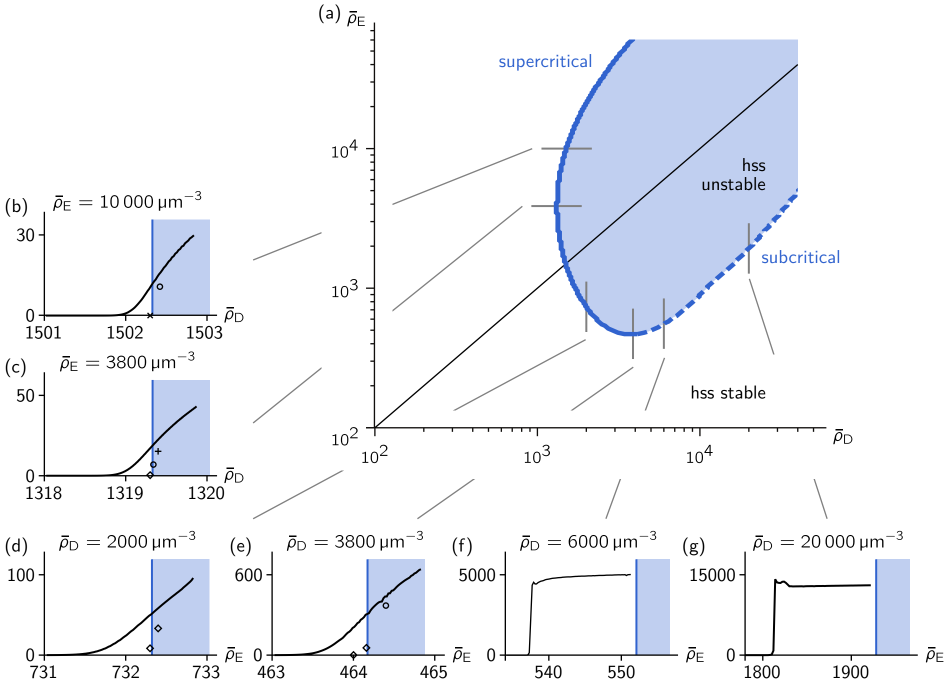

Figure S13. Classification of the onset of pattern formation. (a) Six different points along the onset of linear instability of the homogeneous steady state (hss; dark-blue border of the blue-shaded region) are crossed by tuning the average total MinD or MinE concentrations adiabatically (gray lines). (b–g) The pattern amplitude (of the MinD membrane concentration $m_{\mathrm{d}} + m_{\mathrm{de}}$) is tracked as the total concentration $\bar{\rho}_{\mathrm{D}}$ or $\bar{\rho}_{\mathrm{E}}$ is reduced adiabatically (black line) across the stability boundary of the homogeneous steady state (dark-blue border of the blue-shaded region). For those sweeps where the pattern amplitude decreases to zero continuously (b–e), discrete values of the total concentrations are chosen to perform numerical simulations at fixed average total concentrations for $5 \cdot 10^5$s (plus symbols), $10^6$s (circles), $1.5 \cdot 10^6$s (crosses), or $2 \cdot 10^6$s (diamonds) starting from the concentration pattern determined in the adiabatic sweep for these values of the average total concentrations. The observed super- or subcritical behavior is marked in the concentration phase diagram as a solid and dashed dark-blue border of the instability region, respectively (a). The simulations are performed using COMSOL Multiphysics [18] for the 1+1D model with the parameters given in Table 2. The simulation is initialized with all proteins in the cytosolic species and sinusoidal spatial perturbations with the critical wavenumber $q_{\mathrm{c}}$ at the onset of pattern formation (see Sec. 9.4). The domain length is chosen as $L = 10 \cdot 2\pi/q_{\mathrm{c}}$. For the adiabatic sweeps, weak, constant source terms are introduced at $t = 10^4$s for species $c'_{\mathrm{DD}}$ (b–c) and $c'_{\mathrm{E},1}$ (d–g). In (b–e), the source term is $-2{\cdot}10^{-6}\zeta\mu\mathrm{m}^{-3}\mathrm{s}^{-1}$, in (f) it is chosen as $-2{\cdot}10^{-5}\zeta\mu\mathrm{m}^{-3}\mathrm{s}^{-1}$, and in (g) as $-2{\cdot}10^{-4}\zeta\mu\mathrm{m}^{-3}\mathrm{s}^{-1}$ with the bulk-boundary ratio $\zeta = 0.25\,\mu\mathrm{m}$ (cf. Sec. 8.2).

$\bar{\rho}_{\mathrm{E}}$, respectively. Thereby, the average total concentration is adiabatically varied across the onset of linear instability (see Fig. S13).* For different paths in the concentration phase diagram [see Fig. S13(a)], we track the amplitude of the pattern amplitude while varying the average total concentration [see Figs. S13(b–g)]. As the steady-state pattern is oscillatory (standing or traveling wave), the amplitude is determined as the difference between the maximum and minimum pattern concentrations determined over a time period much larger than the local oscillation period. The setup files for the COMSOL simulation as well as the Mathematica notebook containing the data analysis are available at `https://github.com/henrikweyer/Min-in-vivo` [17].

Close to a supercritical onset of linear instability, the relaxation of the pattern onto the steady state becomes very slow. To determine the pattern amplitude more carefully, discrete values of the average total concentration are chosen, and simulations at fixed average total concentrations are performed starting from the concentration profiles determined in the adiabatic sweep for these concentration values [symbols in Figs. S13(b–e)].

These simulations show that the onset is supercritical for all tested concentrations at the onset of instability if the average total MinD concentration $\bar{\rho}_{\mathrm{D}}$ is low. In contrast, we find multistability and a subcritical onset of pattern formation at the onset of instability at a high average total MinD concentration. We propose that the onset of instability at low $\bar{\rho}_{\mathrm{D}}$ is indeed supercritical for all values of $\bar{\rho}_{\mathrm{E}}$ while the onset at high $\bar{\rho}_{\mathrm{D}}$ is subcritical [solid and dashed blue lines in Fig. S13(a)].

#### 9.4 Quantification of the instability

The initial pattern formation process is dominated by the fastest-growing eigenmode with wavenumber $q_c$ (see Fig. S14). This mode outgrows modes with other wavenumbers due to its larger exponential growth rate. Close to a supercritical onset of pattern formation the wavelength of the final pattern is as well determined by $2\pi/q_c$. The pattern behavior close to such an onset is formalized by the amplitude-equation approach [20, 21]. In the switch model, the onset of pattern formation at small average total concentration of MinD is supercritical (cf. previous section Sec. 9.3). Close to this onset, the final pattern wavelength is well described by the fastest growing mode (see Fig. 4 c). It is an interesting future research direction to analyze this onset in detail and connect the numerical findings with the appropriate amplitude equations for systems with a conservation law and an oscillatory onset of instability [22, 23].

While the real part of the dispersion relation determines the exponential growth or decay of an eigenmode, the imaginary part gives its oscillation frequency. Figure 4 c shows that the period of the standing-wave oscillations in the switch model is again well described by $2\pi/\Im[\sigma(q_c)]$. However, further inside the pattern-forming region, neither the wavelength nor the oscillation period of the final Min patterns can be determined from the properties of the fastest-growing mode. To analyze the pattern

* The data points from the initial period of the simulations are omitted in (b–g) as these only show the initial relaxation of the pattern from the chosen initial condition onto the steady-state pattern.

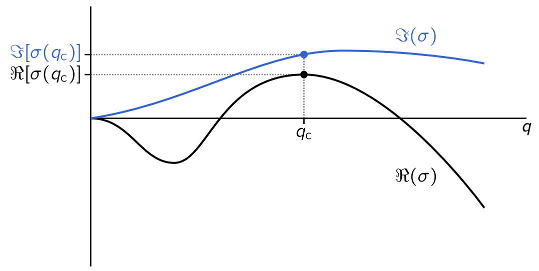

Figure S14. Quantification of the linear instability of the homogeneous steady state. The fastest-growing mode has the wavenumber $q_c$. The real part of the dispersion relation $\Re[\sigma(q_c)]$ determines its growth rate while the imaginary part $\Im[\sigma(q_c)]$ determines the oscillation frequency of the mode.

properties throughout the whole phase diagram, an analysis of the fully developed nonlinear patterns is necessary.

### 10 Parameter study of the switch model reveals standing- and traveling-wave patterns

After the discussion of the region of lateral instability in the $(\bar{\rho}_D, \bar{\rho}_E)$-phase diagram in the last section, we go beyond the linear stability analysis here. In this section, we explore the type of patterns formed by the switch model by numerical simulation of the 1+1D model. The reduced 1+1D model is computationally less expensive and thus allows simulating the concentration phase diagram for different parameter combinations.

To determine a parameter region that not only reproduces the robustness of pattern formation but also the pattern types, we classify the patterns obtained in the numerical simulations and compare the resulting $(\bar{\rho}_D, \bar{\rho}_E)$-phase diagrams for different reaction rates. In the following, we first explain the classification of standing- and traveling-wave patterns. Afterward, the results of the parameter study are presented.

#### 10.1 Pattern classification

For each set of rate parameters and average total concentration values $\bar{\rho}_D, \bar{\rho}_E$ the time evolution of the 1+1D switch model is simulated for 3000 s starting from small random perturbations around the homogeneous steady state. The domain length is $L = 50\,\mu\text{m}$. A notebook with the simulation setup in Mathematica is available under `https://github.com/henrikweyer/Min-in-vivo` [17]. The pattern type obtained at the end of this simulation is classified by analysis of the kymograph of the total MinD membrane concentration $m_d(x,t)+m_{de}(x,t)$ over the last 1000 s. We observe standing- and traveling-wave patterns.

Importantly, the transition from traveling-wave towards standing-wave patterns occurs gradually: The peak concentration of the traveling wave starts oscillating until it decomposes into single oscillating stripes. Moreover, extended standing-wave regimes appear where traveling waves collide with each other or with the domain boundary close to the transition between the pattern types.

Because the transition is gradual, the exact position of the transition line depends on the specific classification criterion. Therefore, in the main text (see Fig. 4 b) the standing- and traveling-wave regions are depicted to gradually transition into each other. This is in agreement with the experimental observation. For simplicity, we will specify a sharp transition here, separating regions of the concentration phase diagrams that show more standing wave-like from those showing more traveling wave-like patterns.

To distinguish traveling-wave patterns from stripe oscillations, we build on the observation that traveling waves moving through the system result in diagonal lines of high surface concentration in the kymograph while stripe oscillations produce isolated spatiotemporal domains of high concentration (see Fig. ??). Consequently, for each oscillation period of length $T$ there is one high-concentration domain expected for a traveling-wave pattern [see Fig. ??(b)]. The total number of high-concentration domains $N$ in a kymograph for a temporal period $\Delta T$ fulfills $N \approx \Delta T/T$.* In contrast, for a standing-wave pattern forming one stripe at midplane, three disconnected high-concentration domains are expected during one oscillation cycle. Shorter wavelengths $\Lambda$ of the standing-wave pattern result in more disconnected domains for each oscillation period [see Fig. ??(a)]. One expects a total number of high-concentration domains $N \approx 2L/\Lambda \, \Delta T/T$. Accordingly, we classify a pattern as a traveling wave if

$$N < (L/\Lambda + 2)\Delta T/T \tag{36}$$

because a domain showing a standing-wave pattern in half the domain and a traveling-wave pattern in the other half fulfills $N \approx (L/\Lambda + 1)\Delta T/T$ if one has $\Delta T/T \gg 1$ and $L/\Lambda \gg 1$.

Instead of showing disconnected high-concentration domains, an (inverted) standing-wave pattern can manifest in disconnected low-concentration domains while the high-concentration domains are connected [see Fig. ??(c)]. This type of standing-wave pattern occurs for average total concentrations very close to the onset of pattern formation at high average total MinD concentrations. Therefore, if the number of high-concentration domains lies below the threshold Eq. (36), we test the same threshold for the number of low-concentration domains [see Fig. ??(b, c)].

To determine the wavelength $\Lambda$, for each time slice of the kymograph the individual structure factor is calculated via the discrete Fourier transform (DFT). The overall structure factor is then determined by averaging the individual structure factors over time. Because the simulation domain has reflective boundaries while the DFT assumes

* The finite temporal length of the kymograph leads to the cutoff of a few domains which increases $N$. Moreover, boundary effects at the ends of the domain lead to a weak oscillation of the concentration maximum of the traveling wave for some parameters, resulting in more than one high-concentration domain for each traveling wave.

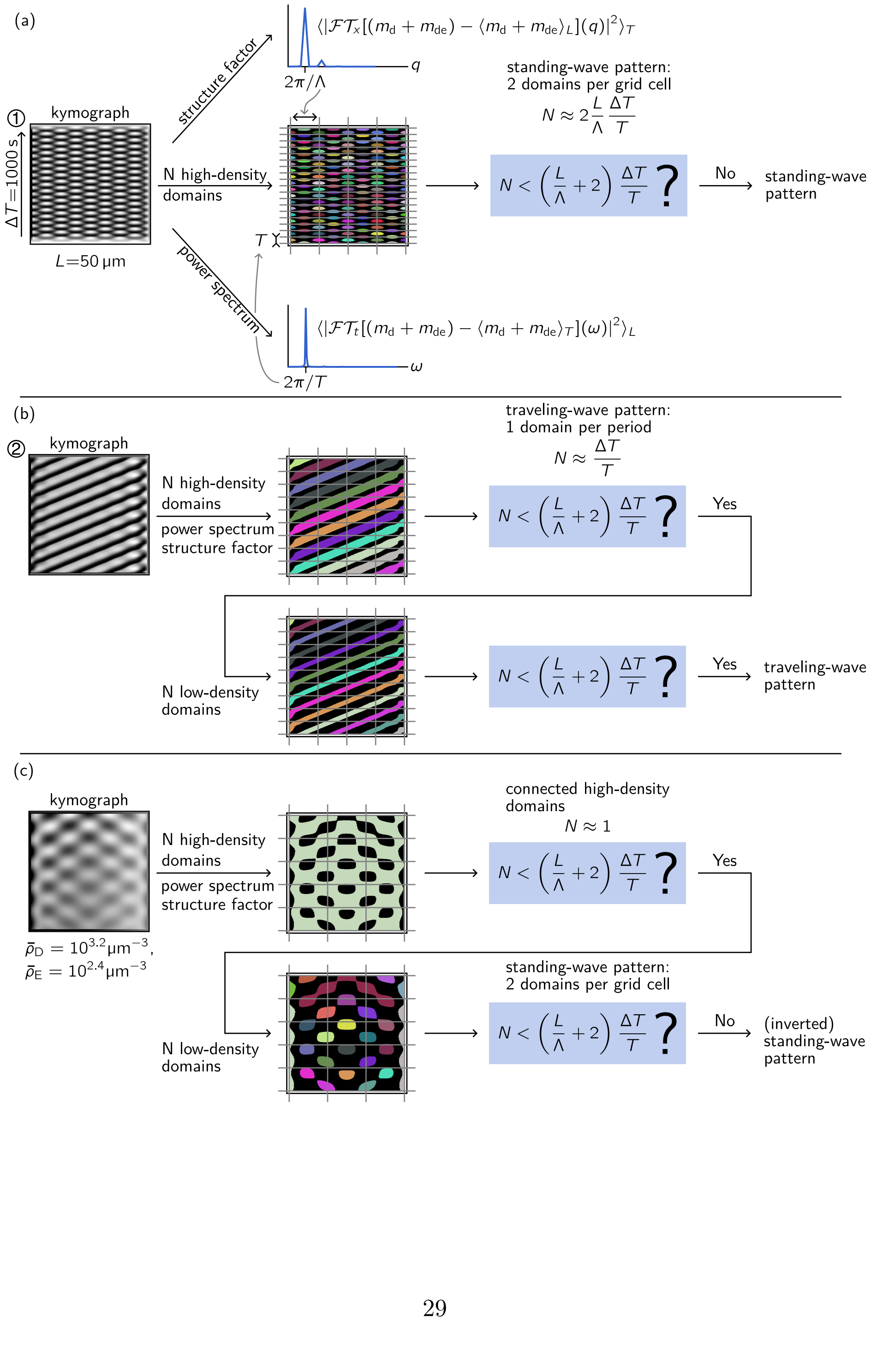
(a)
kymograph
ΔT=1000 s
L=50 µm
structure factor
power spectrum
N high-density domains
⟨|𝓕𝓣x[(md + mde) − ⟨md + mde⟩L](q)|²⟩T
q
2π/Λ
T
⟨|𝓕𝓣t[(md + mde) − ⟨md + mde⟩T](ω)|²⟩L
ω
2π/T
standing-wave pattern:
2 domains per grid cell
N ≈ 2 L/Λ ΔT/T
N < (L/Λ + 2) ΔT/T ?
No
standing-wave pattern
(b)
kymograph
N high-density domains
power spectrum
structure factor
traveling-wave pattern:
1 domain per period
N ≈ ΔT/T
N < (L/Λ + 2) ΔT/T ?
Yes
N low-density domains
N < (L/Λ + 2) ΔT/T ?
Yes
traveling-wave pattern
(c)
kymograph
ρ̄D = 10^3.2 µm^−3,
ρ̄E = 10^2.4 µm^−3
N high-density domains
power spectrum
structure factor
connected high-density domains
N ≈ 1
N < (L/Λ + 2) ΔT/T ?
Yes
N low-density domains
standing-wave pattern:
2 domains per grid cell
N < (L/Λ + 2) ΔT/T ?
No
(inverted) standing-wave pattern

Figure S14. Classification algorithm distinguishing (inverted) standing-wave (a, c) from traveling-wave patterns (b), exemplified for the data points 2 and 1 as well as the simulation with $\bar{\rho}_\mathrm{D} = 2{\cdot}10^{3.2}\mu\mathrm{m}^{-3}, \bar{\rho}_\mathrm{D} = 2{\cdot}10^{2.4}\mu\mathrm{m}^{-3}$ in Fig. 4 b. The wavelength and period of the pattern are determined by discrete Fourier transforms of the kymograph of the MinD membrane concentration $m_\mathrm{d} + m_\mathrm{de}$ (shown in grayscale with white denoting regions of high surface concentration). Then, the kymograph is binarized, and the distinct high-concentration domains are counted. Each distinct high-concentration domain is colored differently. If a standing-wave pattern fills half of the domain while the other half shows a traveling-wave pattern, one expects $N \approx (L/\Lambda + 1)\Delta T/T$ disconnected high-concentration domains if the pattern fulfills $\Delta T/T \gg 1$ and $L/\Lambda \gg 1$. Accordingly, we classify a pattern as (predominantly) standing-wave type if it gives $N \geq (L/\Lambda + 2)\Delta T/T$. To account for standing-wave patterns whose high-concentration domains are connected while the low-concentration domains are disconnected, in a second step the number of the low-concentration domains is tested against the same criterion.

a periodic function, the DFT of each time slice is calculated for the concatenation of the pattern and its reflection. From the structure factor, we determine the dominant Fourier mode $q = 2\pi/\Lambda$ of the pattern as the mode giving the largest value in the structure factor.

Similarly, the power spectrum of the kymograph is calculated to determine the oscillation period $T$. It is computed as the average of the individual power spectra of the time traces at each single spatial point of the pattern which are calculated using the DFT (no concatenation with the reflected time trace). Again the period $T$ is determined from the frequency $\omega = 2\pi/T$ showing the largest value in the power spectrum.

To count the high-concentration domains in the kymograph, the surface concentration is binarized using the standard implementation in Mathematica 13.0 & 13.1 based on cluster variance maximization. From the binarized kymograph, the disconnected high-concentration domains are determined and counted. To count the number of low-concentration domains, the binarized concentrations are inverted before the disconnected domains are counted.

Lastly, if the oscillation period is larger than $\Delta T/2 = 500\,\mathrm{s}$, the pattern is not classified. Moreover, a parameter combination is marked as not showing a pattern if the amplitude in the kymograph is lower than 5% of the minimal pattern membrane concentration:

$$\frac{\max_{\Delta T,L}(m_\mathrm{d} + m_\mathrm{de}) - \min_{\Delta T,L}(m_\mathrm{d} + m_\mathrm{de})}{\min_{\Delta T,L}(m_\mathrm{d} + m_\mathrm{de})} < 0.05\,. \tag{37}$$

In summary, the classification exploits the shape differences of high- and low-concentration domains in the pattern kymographs. This classification only becomes unreliable close to the onset of pattern formation at high average total MinD concentrations because of the long wavelength and period of the patterns. Thus, the kymographs only contain a few pattern domains to compare. Therefore, the inverted standing-wave patterns that occur close to this onset of pattern formation are not clearly distinguished from traveling waves. If less than two periods are contained in the kymograph, the pattern is not classified into either of the two categories.

##### 10.1.1 Pattern classification in the 1+2D model

For the pattern classification in the 1+2D model (see Fig. 4 b), the same classification is used. For the kymograph the membrane concentration $m_{\mathrm{d}} + m_{\mathrm{de}}$ is recorded along the flat membrane of the cylinder (red line in Fig. S10). One has to change the criterion for the stationary state Eq. (37). Due to the reduced local bulk-surface ratio at the cell poles (the spherical caps of the spherocylinder), the stationary state of the system shows inhomogeneous membrane concentrations, even if the system does not undergo a dynamic instability [24]. Therefore, the criterion Eq. (37) is tested for the MinD membrane concentration $m_{\mathrm{d}} + m_{\mathrm{de}}$ only in the middle third of the domain $x \in [L/3, 2L/3]$.

#### 10.2 Parameter study

To find model parameters that reproduce the observed pattern types, our starting points were the study of the skeleton model in *in vivo* geometry performed in Ref. [12] and the linear stability analysis of the switch model for a *in vitro* geometry given in Ref. [13].

Let us first discuss the parameters employed by Denk et al. [13] for the switch model. In their work, the recruitment rates of the reactive and latent MinE states $k_{\mathrm{dE,r}}$ and $k_{\mathrm{dE,l}}$ were tuned over several orders of magnitudes. We choose $k_{\mathrm{dE,r}} = 1\,\mu\mathrm{m}^3\mathrm{s}^{-1}$ and $k_{\mathrm{dE,r}}/k_{\mathrm{dE,l}} = 10^{-4}$ to approximately reproduce the overall region of lateral instability in the concentration phase diagram. In addition, we multiply the hydrolysis rate $k_{\mathrm{de}}$ by 5 for the same reason, resulting in $k_{\mathrm{de}} = 5 \cdot 0.34\mathrm{s}^{-1}$. Moreover, the cytosolic diffusion coefficients measured *in vivo* are used [25] (cf. Table 2). These parameters produce traveling waves throughout almost the whole concentration phase diagram [see Fig. S15(a)]. Only close to the onset of instability at low average total MinD and high MinE concentrations, a standing-wave pattern is observed.

Similarly, we only observe traveling waves when starting from the skeleton model parameters [see Fig. S15(b)]. To use the skeleton model parameters, we extend the parameters from Table 1 by the MinE switch. In order to reproduce the overall instability region, the concentrations are only scaled by a factor of 4 instead of 60 as in Table 1. Moreover, the (scaled) MinE-recruitment rate $k_{\mathrm{dE}} = 0.435/4\mu\mathrm{m}^3\mathrm{s}^{-1}$ is replaced by the rate $k_{\mathrm{dE,r}} = 10k_{\mathrm{dE}}$ for the reactive MinE state and $k_{\mathrm{dE,l}} = 10^{-3}k_{\mathrm{dE}}$ for the latent MinE state. The rate of the conformational switch $\mu = 100\,\mathrm{s}^{-1}$ is chosen as by Denk et al. [13]. In the following, we denote this parameter set as "extended skeleton model".

For the skeleton model, it was argued that the MinD self-recruitment rate has to be high to form pronounced pole-to-pole oscillations in wild-type geometry [12]. We find that this constraint does not carry over to the switch model. Figure S16 shows that the

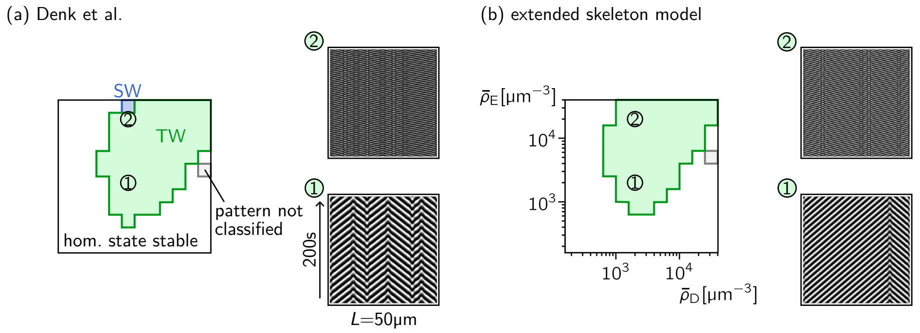

Figure S15. Pattern classification as standing (SW, blue-shaded region) and traveling waves (TW, green-shaded region) in the concentration phase diagrams of the switch model (a) with parameters following Denk et al. [13] and the extended skeleton model (b). In the white region of the phase diagrams the numerical simulation evolves towards the homogeneous stationary state. Patterns not classified are shown as gray-shaded regions (cf. 10.1). (a) The switch model is simulated with the parameters used by Denk et al. [13]. In contrast to the choice of the *in vitro* bulk diffusion coefficients in this work, we choose the cytosolic diffusion coefficients measured *in vivo* (cf. Tables 1, 2). Moreover, the MinE recruitment rates were varied in the study by Denk et al.. We choose $k_{\mathrm{dE,r}} = 1\,\mathrm{\mu m^3 s^{-1}}$ and $k_{\mathrm{dE,r}}/k_{\mathrm{dE,l}} = 10^{-4}$. These values as well as the scaled hydrolysis rate $k_{\mathrm{de}} = 5 \cdot 0.34\mathrm{s}^{-1}$ are employed to approximately reproduce the overall region of lateral instability in the concentration phase diagram. (a) The extended skeleton model describes a parameter set for the switch model which is obtained by extending the parameters of the skeleton model given in Table 1 by rates for a MinE switch. First, the nonlinear rates are scaled by a factor of 4 instead of 60 as in Table 1 in order to move the pattern-forming region to MinD and MinE concentrations comparable with the experimental phase diagram. Second, the (scaled) MinE-recruitment rate $k_{\mathrm{dE}} = 0.435/4\mathrm{\mu m^3 s^{-1}}$ is split into the rate $k_{\mathrm{dE,r}} = 10 k_{\mathrm{dE}}$ for the reactive MinE state and $k_{\mathrm{dE,l}} = 10^{-3} k_{\mathrm{dE}}$ for the latent MinE state. For both parameter sets example kymographs of traveling wave patterns for two different concentration combinations are shown. The MinD membrane concentration $m_{\mathrm{d}} + m_{\mathrm{de}}$ is shown in grayscale with white denoting high membrane concentration.

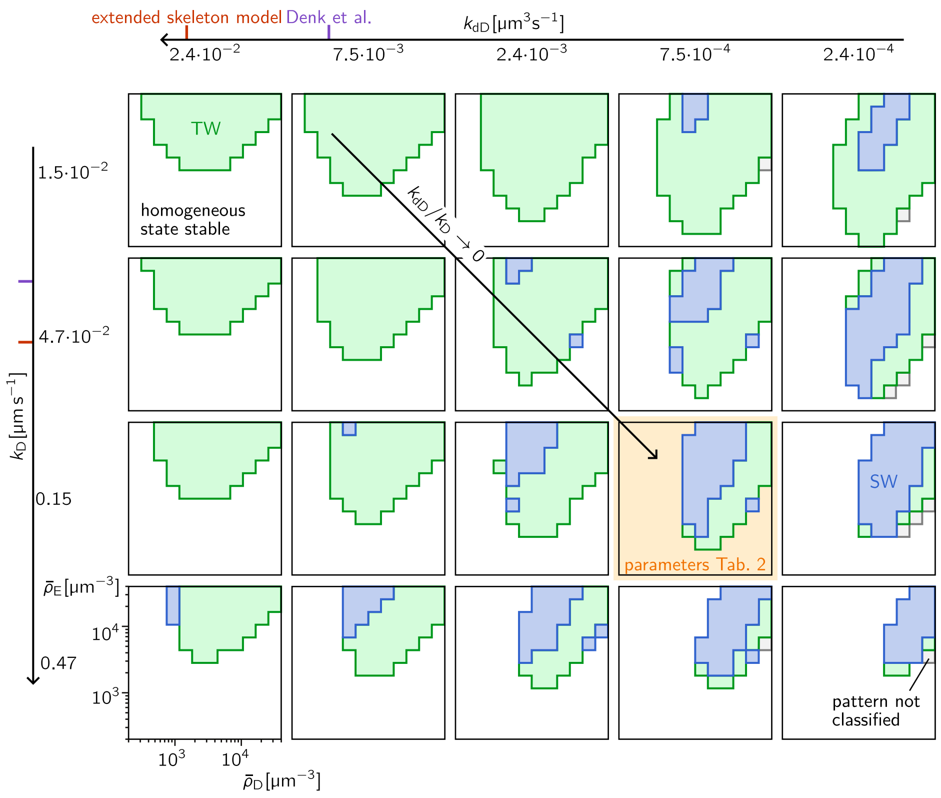

Figure S16. The effect of MinD attachment rates $k_{\mathrm{D}}$ and $k_{\mathrm{dD}}$ on the pattern types in the concentration phase diagram. For each combination of values for the rates $k_{\mathrm{D}}$ and $k_{\mathrm{dD}}$, the 1+1D model is simulated for different combinations of average total concentrations $\bar{\rho}_{\mathrm{D}}$ and $\bar{\rho}_{\mathrm{E}}$. For each combination of concentrations, the pattern type is determined as explained in Sec. 10.1. The result is a concentration phase diagram for each rate combination showing the region of standing waves (blue-shaded), traveling waves (green-shaded), and patterns not classified (gray-shaded). For the other combinations of average total concentrations, the homogeneous state is stable. The phase diagram corresponding to the final parameter set given in Table 2 is shaded in orange. The values of $k_{\mathrm{D}}$ and $k_{\mathrm{dD}}$ used by Denk et al. and in the extended skeleton model are marked on the parameter axes in purple and red, respectively. The parameters kept constant are set according to Table 2.

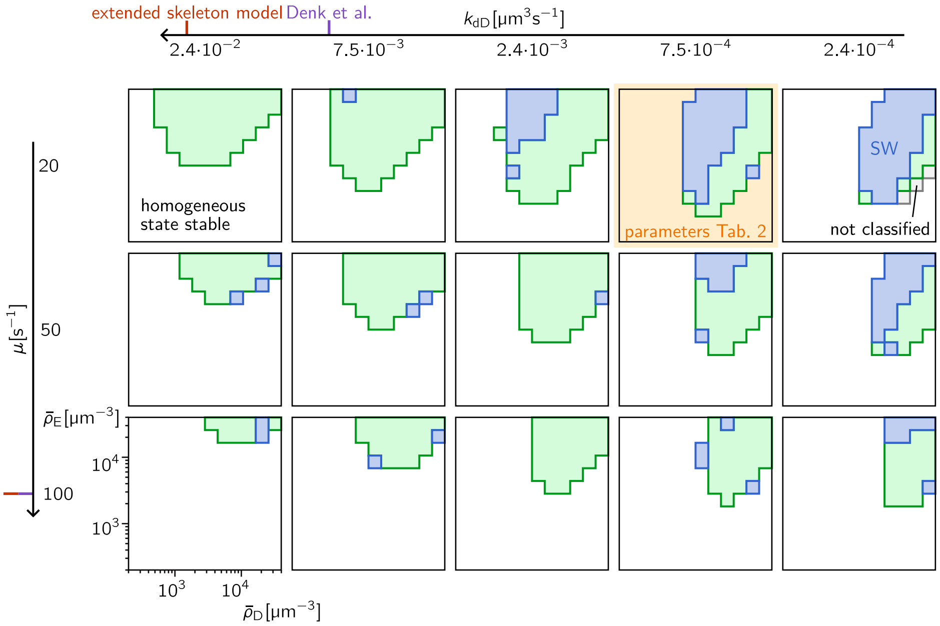

Figure S17. The effect of the rate of the conformational switch on the pattern types in the concentration phase diagram. The color code is the same as in Fig. S16. Again, the parameters not varied are given in Table 2.

switch model only forms standing-wave patterns (giving rise to pole-to-pole oscillations in short cells of wild-type length) in large regions of the phase diagram if the MinD self-recruitment rate $k_{dD}$ is sufficiently low compared to the rate of spontaneous MinD membrane attachment $k_D$. Consequently, the MinE switch allows for standing-wave patterns already if the MinD self-recruitment is weak. The values of the parameters $k_D$ and $k_{dD}$ used by Denk et al. and in the extended skeleton model are marked in Fig. S16 for comparison. Note however that the other parameters not varied are fixed to the values given in Table 2 different from the values used by Denk et al. and in the extended skeleton model.

Moreover, Figure S17 shows that a reduced rate of the conformational switch $\mu$—although still larger than all other rates—also enlarges the standing-wave region in the concentration phase diagram.

Taken together, this selection of the parameter sweeps explains the most important parameter changes that lead to the final parameter set given in Table 2: The nonlinear MinD attachment, i.e., the MinD self-recruitment is strongly reduced as well as the rate of the conformational MinE switch. It will be an interesting task for future research to uncover the mechanistic processes underlying standing-wave and traveling-wave formation based on the overview of the parameter space we give here.

##### 10.2.1 Wavelength and period of the nonlinear patterns

Finally, after matching the region of pattern formation as well as the observed pattern types, the third layer of detail is the quantitative comparison of the pattern wavelength and period.

The analysis of the pattern wavelength for different rate parameters shows that it is almost constant throughout the concentration phase diagram (see Fig. S18). The parameter study shows that the wavelength is strongly influenced by the self-recruitment rate of MinD $k_{\mathrm{dD}}$. This behavior is similar to the observation in the skeleton model that the self-recruitment rate sets the position where the new polar zone starts to grow [12]. Thus, the value of $k_{\mathrm{dD}}$ is chosen to reproduce the experimentally determined wavelength. The value of the linear attachment rate $k_{\mathrm{D}}$ then follows from the matching of the pattern types.

While reproducing the experimental wavelength, also the oscillation period should fit the experiment. The length and time scales of the system are connected via the diffusion coefficients, and it is not obvious that both can be tuned independently as the values of the cytosolic diffusion coefficients are fixed to their experimentally measured values. We find in the switch model that the hydrolysis rate $k_{\mathrm{de}}$ tunes the oscillation period (see Fig. S19) while it does not strongly affect the pattern wavelength. The value chosen for $k_{\mathrm{de}}$ is determined such that the region of pattern formation, as well as the oscillation period, is reproduced.

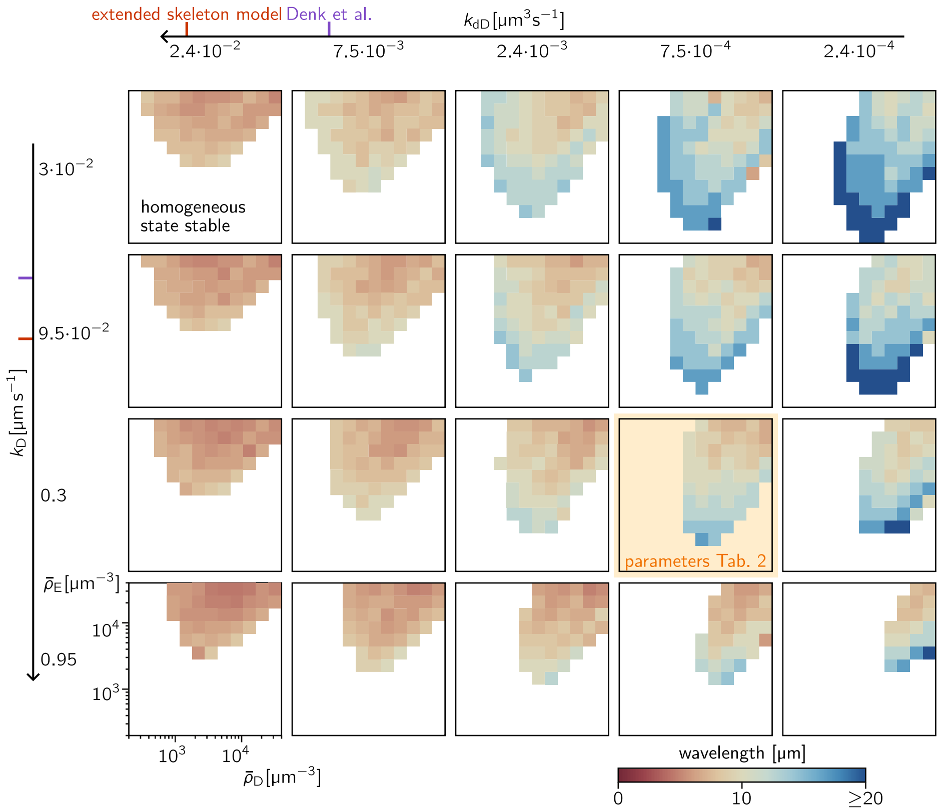

Figure S18. Dependence of the pattern wavelength on the linear attachment and self-recruitment rates $k_D$, $k_{dD}$ of MinD. The fill color denotes the observed pattern wavelength according to the color legend (bottom right). An increasing self-recruitment rate $k_{dD}$ decreases the observed pattern wavelength. The phase diagram corresponding to the chosen parameter set (cf. Table 2) is marked in orange. Again, the parameters not varied are given in Table 2.

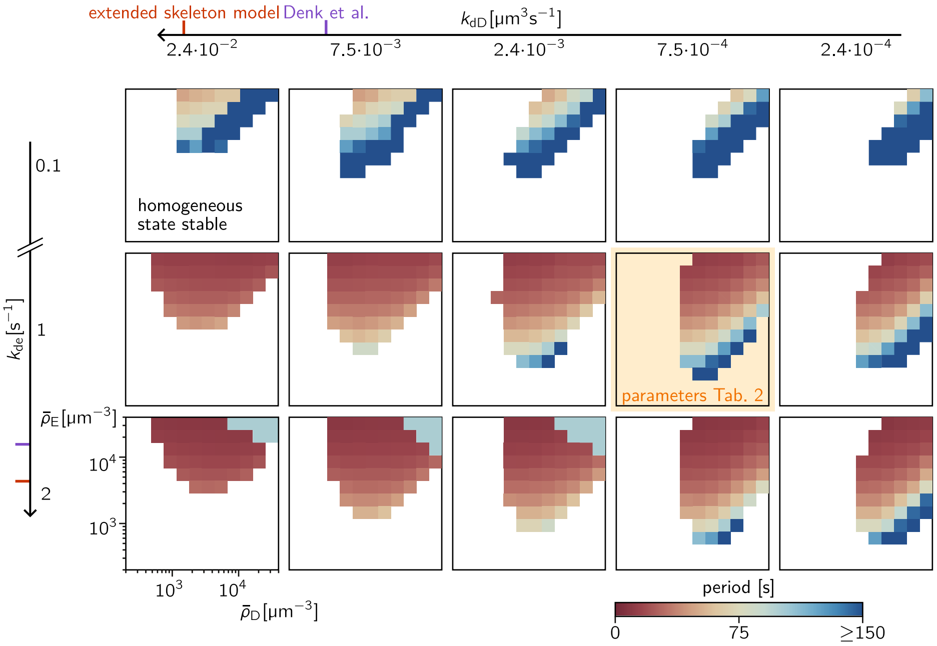

Figure S19. Dependence of the local oscillation period on the hydrolysis rate $k_{de}$ and the MinD self-recruitment rate $k_{dD}$. The fill color denotes the oscillation period according to the color legend (bottom right). The hydrolysis rate influences both the onset of pattern formation at low average total MinE concentrations and the oscillation period. At the high hydrolysis rate, $k_{de} = 2\,s^{-1}$ and high average total MinD and MinE concentrations an abrupt change in the pattern oscillation period can be observed. Again, the parameters not varied are given in Table 2.
